# Dopamine release at the time of a predicted aversive outcome causally controls the trajectory and expression of conditioned behavior

**DOI:** 10.1101/2022.04.15.488530

**Authors:** Munir Gunes Kutlu, Jennifer Tat, Jennifer E. Zachry, Erin S. Calipari

**Author notes:** **Corresponding Author Erin S. Calipari, PhD**, Assistant Professor, Department of Pharmacology, Department of Molecular Physiology and Biophysics, Department of Psychiatry and Behavioral Sciences, Vanderbilt Center for Addiction Research, Vanderbilt Brain Institute, Vanderbilt University 865F Light Hall, 2215 Garland Avenue, Nashville, TN 37232.

## Abstract

**Background:** The inability to predict when aversive stimuli will and will not occur in is a hallmark of anxiety and stress disorders. Dopamine release in the nucleus accumbens (NAc) is sufficient and necessary for aversive learning and has been linked to both anxiety and stress disorder symptomatology. Thus, understanding how dopamine controls associative learning in response to aversive stimuli is critical to understanding the role of dopamine in behavior in health and disease.

**Methods:** Using an optical dopamine sensor combined with *in-vivo* fiber photometry in the NAc core of male and female C57BL/6J mice (N=38), we recorded dopamine responses to expected and omitted aversive outcomes during learning. We derived predictions from a theory-driven model of associative learning (Kutlu-Calipari-Schmajuk, KCS model) and tested the causality of these predictions using optogenetics.

**Results:** Dopamine release was evoked by the predicted omission of aversive stimuli in a fashion that cannot be explained by dopamine as a reward-based prediction signal. The magnitude of the dopamine response during omissions scaled with predictions about the probability of their occurrence; however, dopamine did not track the associative value of predictive cues. Finally, we showed that the observed effects are causal to learned behavior and can only be explained by dopamine signaling the perceived saliency of predicted aversive events.

**Conclusions:** We elucidate the role of NAc core dopamine signaling in aversive learning in a theory-based and stimulus-specific fashion and offer potential avenues for understanding the neural mechanisms involved in anxiety and stress disorders.

## Introduction

The execution of adaptive behavior depends on the ability of organisms to predict potential threats in their environment. To this end, animals learn to predict when aversive stimuli will occur and learn what actions are necessary to avoid contexts and situations with potential negative outcomes. However, equally important, is the ability to learn when aversive stimuli are *not* likely to be presented. This balance allows animals to avoid potential negative outcomes while still exploring their environment when it is safe and unlikely to result in harm(1,2). The inability to effectively discriminate between these situations and update information about these relationships is a hallmark of pathological disease states, such as PTSD and anxiety disorders(3–5). Thus, understanding the neural basis of these behaviors and how neuromodulatory systems in the brain control them is critically important to understanding this behavior in health and conceptualizing what dysfunction in the neural systems that control this means for disease states.

Dopamine is widely studied in the context of associative learning(6–12) as well as clinically in anxiety and stress disorders where its dysregulation is a disease hallmark(13–15). While historical work has linked dopamine to reward processing and motivation, dopamine release in the nucleus accumbens (NAc) is sufficient and necessary for the acquisition and expression of conditioned associations for both rewarding and aversive stimuli(10,11,16–19). Work has more recently begun to study the role of rapid dopamine signaling in the processing of aversive stimuli; however, there are competing ideas about how dopamine release in striatal regions - such as the NAc core, which we focus on in this study - causally controls the expression of associated conditioned behavioral responses. Some have suggested that the dopamine release observed at the time of an omitted aversive outcome signals relief/safety and functions as a reward-based prediction signal(10,11,20). Others, including our group, have hypothesized that dopamine acts to transmit a saliency signal that helps to update adaptive behavior regardless of the valence of the stimulus or outcome(17,21–25). Differentiating these hypotheses is particularly critical as these accounts would hypothesize opposite effects of dopamine in updating behavior associated with aversive stimuli. Similarly, how to conceptualize dopaminergic dysfunction observed in patients suffering from stress disorders would be different and could influence how treatment is approached.

Here we combine dopamine recordings and computational modeling in mice to outline how dopamine contributes to conditioned behavioral responses for aversive stimuli. We define how dopamine release guides the ability of an animal to discriminate between situations where aversive outcomes are or are not likely to occur. To this end, we combine *in vivo* approaches for optical recording and manipulations of dopamine terminals in the NAc with behavioral tasks where shock presentation and omission is predicted by a cue. These data show that dopamine in the NAc plays a causal role in guiding animals to learn to predict when aversive stimuli will and will not be presented in an environment. However, this occurs through the modulation of dopamine release at the time of the predicted outcome (shock or omission), not at the time of the cue as other reward/prediction-based accounts of dopamine in learning and memory have hypothesized. We then combine this with computational modeling approaches to demonstrate that these patterns can only be explained by perceived saliency coding, rather than associative or valence-based variables. Our results demonstrate that NAc core dopamine release plays a role in aversive learning in a way that is more complex than previously hypothesized and may be an important potential target for anxiety and stress disorders where impaired fear/safety learning is a critical component.

## Materials and Methods

### Animals

Male and female 6-to 8-week-old C57BL/6J mice (N=38) obtained from Jackson Laboratories (Bar Harbor, ME; SN: 000664) were kept 5 per cage and maintained on a 12-hour reverse light/dark cycle, with all behavioral testing taking place during the light cycle. Animals were given *ad libitum* access to food and water. All experiments were conducted in accordance with the guidelines of the Institutional Animal Care and Use Committee (IACUC) at Vanderbilt University School of Medicine. Experimenters were blind to experimental groups throughout behavioral experiments.

### Surgical Procedure

At least 1 hour prior to surgery, mice were administered Ketoprofen (5 mg/kg) via subcutaneous injection. Animals were anesthetized using isoflurane (5% for induction and 2% for maintenance) and placed on a stereotaxic frame (David Kopf Instruments). Ophthalmic ointment was applied to the eyes throughout surgical procedures. A midline incision was then made down the scalp and a craniotomy was performed with a dental drill using aseptic technique. Using a .10-mL NanoFil syringe (WPI) with a 34-gauge needle, AVV5.CAG.dLight1.1 (UC Irvine;(26)) was unilaterally infused into the NAc (bregma coordinates: anterior/posterior, +1.4 mm; medial/lateral, +1.5 mm; dorsal/ventral, −4.3 mm; 10° angle) at a rate of 50 nL/min for a total volume of 500 nL. Following infusion, the needle was kept at the injection site for seven minutes before being slowly withdrawn. Fiber-optic cannulas (400 μm core diameter; .48 NA; Doric) were then implanted in the NAc and positioned immediately dorsal to the viral injection site (bregma coordinates: anterior/posterior, + 1.4 mm; medial/lateral, + 1.5 mm; dorsal/ventral, −4.2 mm; 10° angle) before being permanently affixed to the skull using adhesive cement (C&B Metabond; Parkell). Follow-up care was performed according to IACUC/OAWA and DAC standard protocol. Animals were allowed a minimum of six weeks to recover in order to ensure efficient viral expression before commencing experiments. For the optogenetic experiments, AAV5.Ef1a.DIO.hChR2.eYFP (ChR2; UNC vector core), Chrimson.FLEX: AAV5-Syn-FLEX-rc[ChrimsonR-tdTomato] (Chrimson; Addgene) or AAV5-Ef1a-DIO.eNpHR.3.0-eYFP (NpHR; Addgene) and AAV9.rTH.PI.Cre.SV40 (Addgene;(27)) were injected into the VTA (unilaterally for ChR2 and Chrimson and bilaterally for NpHR; bregma coordinates: anterior/posterior, −3.16 mm; medial/lateral, +0.5 mm; dorsal/ventral, −4.8 mm) of C57BL/6J mice. Unilateral (for ChR2 and Chrimson) or bilateral (for NpHR) 200 μm core diameter fiber optic implants were placed into the NAc core (bregma coordinates: anterior/posterior, +/-1.4 mm; medial/lateral, +1.5 mm; dorsal/ventral, −4.3 mm; 10° angle; at a rate of 50 nL/min for a total volume of 500 nL). This allowed for the photostimulation or photoinhibition of dopamine response only in dopamine terminals that project from the VTA and synapse in the NAc core. Control animals received AAV5.Ef1a.DIO.eYFP injections into the VTA instead of ChR2 or NpHR.

### Fiber Photometry

The fiber photometry system used two light-emitting diodes (490 nm and 405 nm; Thorlabs) controlled by an LED driver (Thorlabs). The 490 nm light source was filtered with a 470 nm (the excitation peak of dLight1.1) bandpass filter and the 405nm light source was used as an isosbestic control(26). Light was passed through an optical fiber (400 μm, .48 NA; Doric) that was coupled to a chronically implanted fiber optic cannula in each mouse. LEDs were controlled via a real-time signal processor (RZ5P; Tucker-Davis Technologies) and emission signals from each LED were determined by multiplexing. Synapse software (Tucker-Davis Technologies) was used to control the timing and intensity of the LEDs and to record the emitted fluorescent signals upon detection by a photoreceiver (Newport Visible Femtowatt Photoreceiver Module; Doric). LED power (125 μW) was measured daily and maintained across trials and experiments. For each event of interest (e.g., cue presentation, footshock), transistor-transistor logic (TTL) signals were used to timestamp onset times from Med-PC V software (Med Associates Inc.) and were detected via the RZ5P in the Synapse software (see below). A built-in low-pass filter on the Synapse software was set to 10 Hz to eliminate noise in the fiber photometry raw data.

### Behavioral Experiments

#### Aversive conditioning

<u>Initial Training.</u> We employed an aversive conditioned inhibition design where mice received two consecutive training sessions (Sessions 1 - 2) where mice were presented with six trials of house light presentations (Fear cue; 10 seconds in duration) followed by a footshock (1 mA, 0.5 second duration) and 12 trials of a combination of house light and white noise presented concurrently with no shock presented (Fear+Omission cue; 85 dB; both 10 seconds in duration) with a variable inter-trial interval (35 - 55 seconds). The session was organized where mice received one Fear cue (house light) trial followed by two Fear+Omission (houselight and whitenoise) trials, which repeated until the total number of each trial type was reached. <u>Test Session.</u> During a subsequent test session, mice received three Fear cue (house light only) and three Fear+Omission cue trials (house light and white noise). During these sessions, no footshock was presented for either of these trial types. Freezing response was measured during both training and test sessions. The freezing response was defined as the time (seconds) that mice were immobile (lack of any movement including sniffing) during the tone period and calculated as percentage of total cue time. We converted freezing durations to percentages of total cue time ((freezing time * 100)/ stimulus duration). Lower freezing levels during the Fear+Omission cue trials compared to the Fear cue only trials at the testing session were considered as successful learning of the task contingencies.

#### Optogenetic photostimulation

In a group of C57BL/6J mice, AAV5.Ef1a.DIO.hChR2.eYFP (ChR2; UNC vector core) and AAV9.rTH.PI.Cre.SV40 (Addgene) were injected into the VTA and a 200 μm fiber optic implant was placed into the NAc core. This allowed for photostimulation of dopamine response only in dopamine terminals that project from the VTA to NAc core. Control animals received AAV5.Ef1a.DIO.eYFP injections into the VTA instead of ChR2. Mice received the same training outlined above. During the two initial training sessions (Sessions 1-2), blue laser stimulation (470 nm, 1 second, 20 Hz, 8 mW) was delivered into the NAc core at the offset of the Fear+Omission cue trials at the time of the predicted, but absent footshock for 1s. In the following session (Session 3), mice received the same testing sessions as described above; however, there was no laser stimulation during this testing session. Freezing was measured during both training and testing sessions.

In a separate group of C57BL/6J mice, AAV5-Ef1a-DIO.eNpHR.3.0-eYFP (NpHR; Addgene) and AAV9.rTH.PI.Cre.SV40 (Addgene;(27)) were injected into the VTA and a 200 μm fiber optic implant was placed into the NAc core. These mice received the same training and stimulation treatments as the ChR2 mice; however, during each training session, we delivered yellow laser stimulation (590 nm, 1 second, constant, 8 mW) into the NAc core at the offset of the Fear+Omission cue trials at the time of footshock omission for 1 second. In the next session, mice received the same training test without laser stimulation as was done in the ChR2 mice. Freezing was measured during both training and testing sessions.

#### Optogenetic photostimulation/inhibition combined with concurrent optical dopamine recording

*<u>Photostimulation</u>*. A group of C57BL/6J mice was injected with Chrimson.FLEX: AAV5-Syn-FLEX-rc[ChrimsonR-tdTomato] (Chrimson; Addgene) and AAV9.rTH.PI.Cre.SV40 (Addgene) into the VTA, and AVV5.CAG.dLight1.1 (UC Irvine) was injected into the NAc core as described above. A 400 μm fiber optic was implanted into the NAc core. Using the same paradigm described above yellow laser stimulation (590 nm, 1 second, constant, 8 mW) was delivered into the NAc core at the offset of the Fear+Omission cue while recording fluorescent signals emitted from dLight1.1 in the same animal using fiber photometry. We have previously validated this approach(17). *<u>Photoinhibition</u>*. In a separate group of C57BL/6J mice, an inhibitory opsin (AAV5.hSyn.eNpHR3.0.mCherry) was injected bilaterally into the VTA and AVV5.CAG.dLight1.1 (UC Irvine) was injected into the NAc core as described above. Bilateral 400 μm fiber optic implants were placed into the NAc core. Using a within subject design, during a single training session, mice received 5 tone (2.5 kHz, 85 dB) or 5 houselight presentations (intermixed randomly, both 5 seconds in duration), both paired with a footshock outcome (1 mA, 0.5 second duration). Dopamine terminals were inhibited during the footshock following one cue (Inhibited cue; either the tone or the houselight counterbalanced) via yellow laser stimulation (590 nm, 1 second, constant, 8 mW). The footshock following the other cue was not paired with any laser stimulation (Non-inhibited cue). In the following session, mice received 5-8 presentations of the Inhibited and Non-inhibited cues and freezing response and dopamine responses were measured in response to each cue. Dopamine responses were recorded unilaterally via fiber photometry during the expected but absent footshock in each case.

#### Computational simulations using the KCS Model

Using a theory-driven neural network model of associative learning, the Kutlu-Calipari-Schmajuk (KCS) model(17), we simulated the conditioned response during the conditioned inhibition paradigm. One advantage of this approach is that we are able to test hypotheses regarding what happens to conditioned response when a signal is artificially enhanced or suppressed. Here, we utilized this approach to examine whether the dopamine response to the expected but absent footshock represents a perceived saliency signal by simulating optogenetic stimulation and inhibition of the dopamine signal. For each KCS model simulation (see supplementary methods for the details of the model), we determined 6 free parameters (ITI duration, cue duration, outcome duration, cue value, outcome value, and number of training trials) to mimic the experimental design of the behavioral experiments. Although these values are chosen arbitrarily, we kept them constant throughout the study. For all simulations we used ITI duration of 200 t.u. and cue and outcome values of 1. For the conditioned inhibition (safety learning simulations), we used a total of 18 CS1-outcome trials intermixed with 36 CS1+CS2-no outcome trials. For the inhibitory optogenetic manipulations, we assumed that the perceived saliency (or prediction error) of the expected but absent outcome during the CS1+CS2-no outcome trials was −1, and for the excitatory optogenetics simulations, we set that value to 2 in order to mimic dopamine response. For the eYFP control simulations, we let the model determine the perceived saliency (or prediction error) of the outcome. We printed out the simulated conditioned response values at the offset of the cue presentations (t.u. 30). For the inhibited versus non-inhibited outcome simulations, we used the same model parameters and simulated 5 cue-outcome trials followed by 5 cue-no outcome trials. During the initial 5 cue-outcome simulations, the perceived saliency of the outcome was set to −1 to mimic optogenetic inhibition of dopamine. For the simulations of the non-inhibited cue, we let the model determine the perceived saliency of the outcome. We printed out the conditioned response as well as perceived saliency, associative strength (V_CS-US_), and prediction error values at the offset of the test cue presentations.

For a more detailed explanation of the KCS model as well as several other methodological details see the supplementary methods.

## Results

### Dopamine release in the NAc core is evoked by the omission of a predicted shock; this response scales positively with the strength of prediction

Our first goal was to determine how NAc core dopamine release tracked the predicted presentation (or omission) of an aversive stimulus (**Figure 1a-c**; **Figure S1a-b**). To this end, animals were trained in a conditioned inhibition paradigm where they learned to predict when shock would and would not be likely to be presented based on learned antecedent cues. During each session, mice were exposed to two trial/cue types: 1) a Fear cue (house light) that predicted the delivery of a footshock (**Figure 1d**) or 2) a compound cue (whitenoise and house light) that signaled that no shock would be presented at the end of the cue period (termed Fear+Omission cue throughout the manuscript). Learning on this task is characterized by freezing less to the Fear+Omission cue than the Fear cue alone since the added Omission cue is predictive of the absence of shock. This reduced behavioral response in the presence of the inhibitory cue (Omission cue) is traditionally called conditioned inhibition. Indeed, mice showed robust learning (exhibiting conditioned inhibition) following 2 sessions of training indicated by weaker freezing response to the Fear+Omission cue compared to the Fear cue alone during testing (**Figure 1e-f**; **Figure S2**).

**Figure 1.**
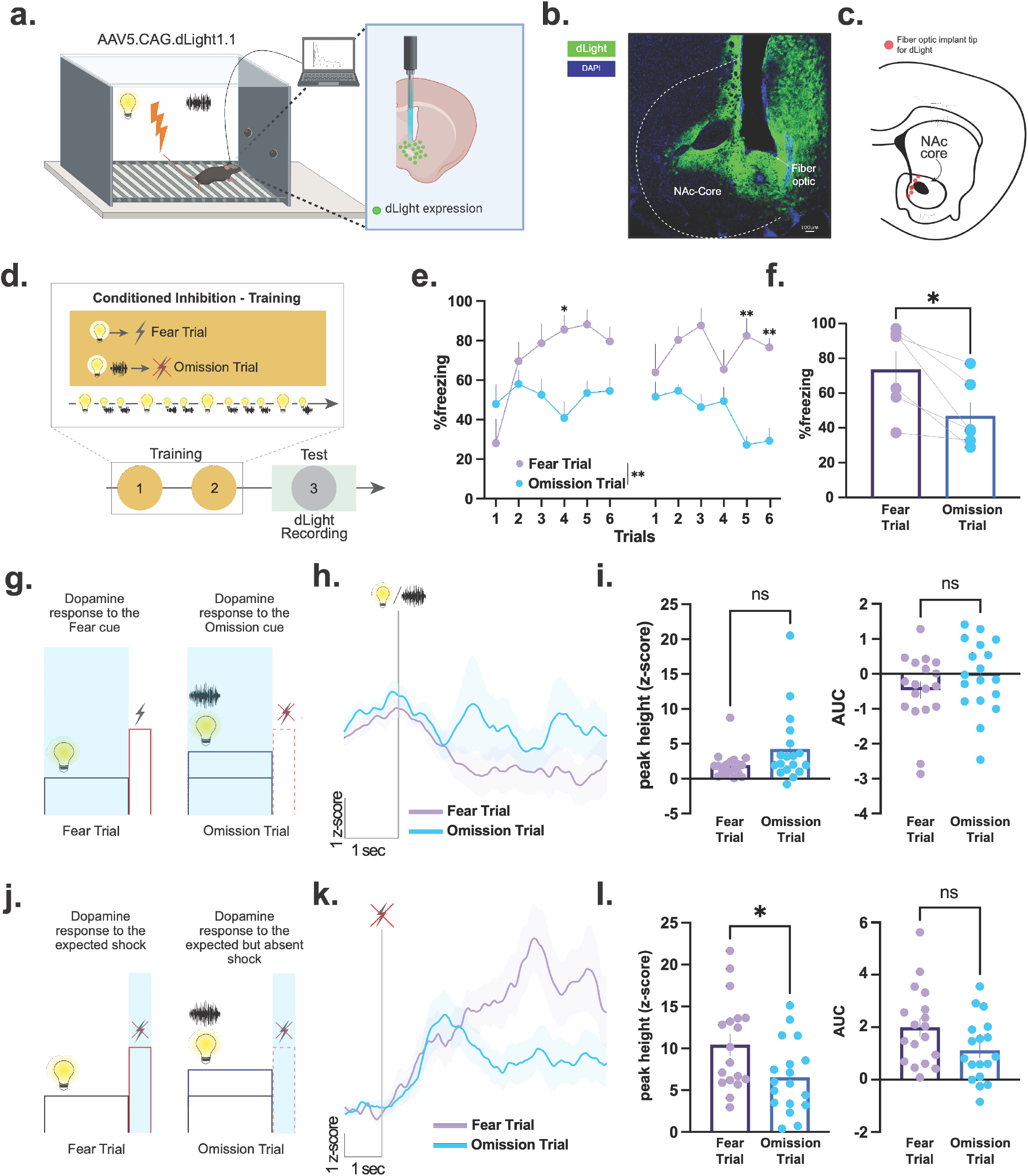
NAc core dopamine release is evoked by the omission of predicted aversive outcomes and scales with predictions. **(a)** Mice (n=6) received unilateral injections of the fluorescent dopamine sensor dLight1.1 in the nucleus accumbens (NAc). **(b)** A fiber optic cannula was placed directly above the injection site in the NAc core. Representative histology showing viral expression (green) restricted to the NAc core and **(c)** schematic showing fiber optic placements (red) in experimental animals. **(d)** Aversive conditioned inhibition paradigm. During training there were two trial types: 1). the Fear cue (houselight) was paired with a footshock and 2). the Fear+Omission cue was a compound cue (house light and white noise together) that signaled that the footshock would not be presented. **(e)** Trial-by-trial freezing responses to the Fear cue and Fear+Omission cue during training (RM ANOVA main effect of group F_(1, 10)_= 19.87 p=0.025, n=6) and **(f)** averaged freezing responses during testing (paired t-test, *t*_5_=3.53, *p*=0.0150, n=6 mice). Mice exhibited robust conditioned inhibition learning – i.e. stronger freezing to the Fear cue alone compared to the Fear+Omission cue. **(g-h)** Dopamine response at the time of the Fear cue as compared to the Fear+Omission cue did not differ. **(i)** Peak height (Nested t-test, *t*_10_=1.56, *p*=0.1480, n=6 mice) and area under the curve (AUC; Nested t-test, *t*_10_=0.95, *p*=0.3618, n=6 mice) of the dopamine response to the Fear cue and Fear+Omission cue. **(j-k)** Dopamine response at the time of the expected but omitted footshock following the Fear cue was stronger compared to the dopamine response to the omitted footshock after the Fear+Omission cue. **(l)** Both peak height (Nested t-test, *t*_34_=2.39, *p*=0.0221, n=6 mice) and area under the curve (AUC; Nested t-test, *t*_10_=1.77, *p*=0.1058, n=6 mice) values. Data represented as mean ± S.E.M. * *p* < 0.05, ** *p* < 0.01; ns = not significant.

**Figure 2.**
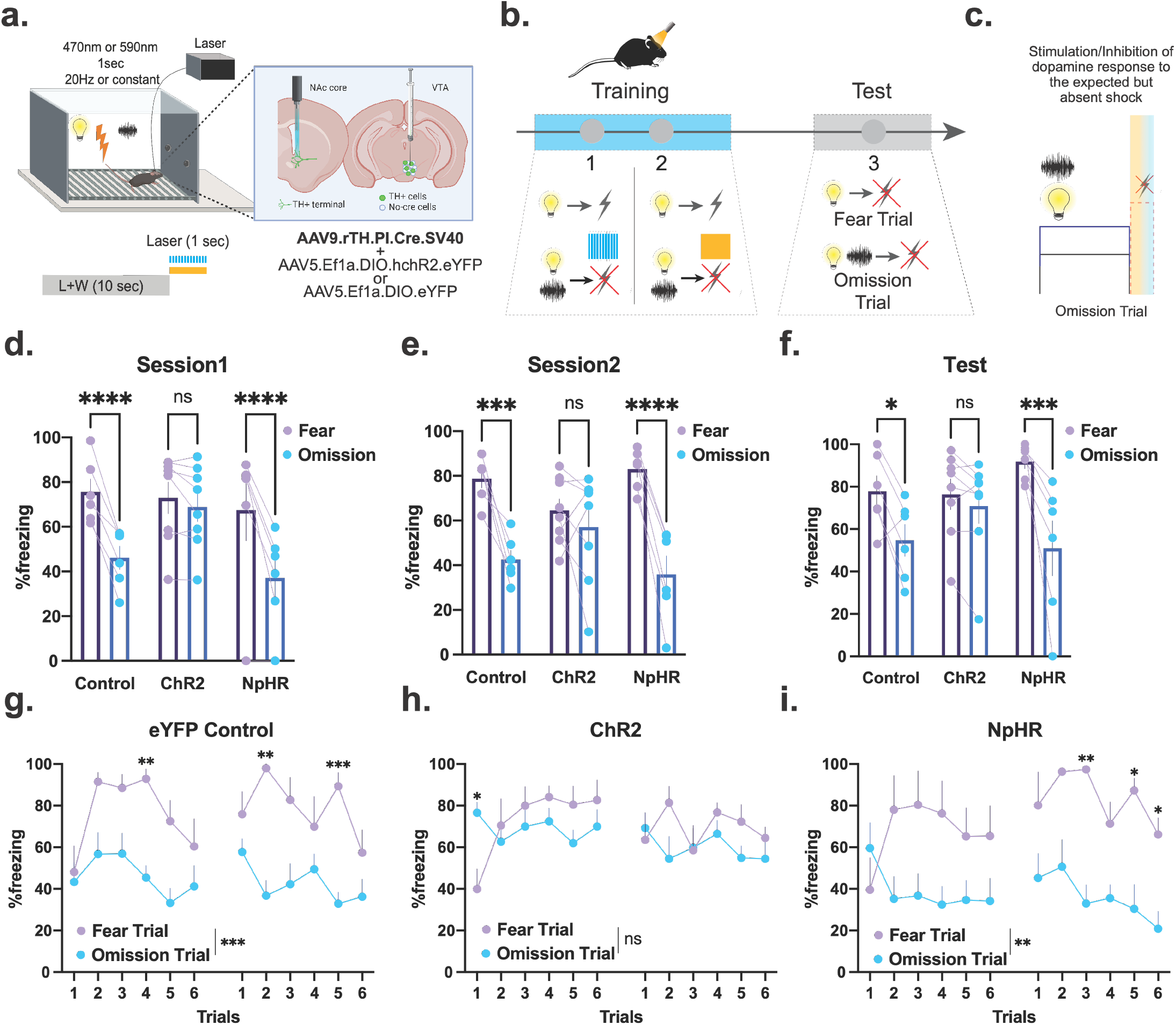
Optogenetic stimulation of the NAc core dopamine terminals at the time of the expected but absent footshock disrupts conditioned inhibition learning. **(a)** AAV5.Ef1a.DIO.eYFP (eYFP), AAV5-Ef1a-DIO.eNpHR.3.0-eYFP (NpHR), or AAV5.Ef1a.DIO.hchR2.eYFP (ChR2) were co-injected with AAV9.rTH.PI.Cre into the VTA to achieve dopamine-specific expression of excitatory or inhibitory opsins. (**b-c**) Dopamine release was optogenetically evoked (via blue laser – ChR2) or inhibited (via yellow laser + NpHR) from terminals in the NAc core at the time of the omitted shock following the Fear+Omission cue (houselight+whitenoise). Averaged freezing responses to the Fear cue (houselight) and to the Fear+Omission cue following optogenetic manipulations during **(d)** Session 1 (RM ANOVA Group x Cue interaction *F*_(2,17)_= 11.19, *p*=0.0008; eYFP Control Sidak post-hoc p<0.0001; ChR2 Sidak post-hoc p>0.05; NpHR Sidak post-hoc p<0.0001; n=20) and **(e)** Session 2 (RM ANOVA Group x Cue *F*_(2,17)_= 8.27, *p*=0.0031; eYFP Control Sidak post-hoc p<0.01; ChR2 Sidak post-hoc p>0.05; NpHR Sidak post-hoc p<0.0001; n=20) of training and **(f)** testing (RM ANOVA Group x Cue interaction *F*_(2,17)_= 4.97, *p*=0.0199; eYFP Control Sidak post-hoc p<0.05; ChR2 Sidak post-hoc p>0.05; NpHR Sidak post-hoc p<0.001; n=20). **(g)** Control mice showed robust learning indicated by stronger freezing response to the Fear cue alone compared to the Fear+Omission cue during training (RM ANOVA main effect of cue type F_(1, 10)_= 33.00 p=0.0002, n=20). **(h)** Optogenetic stimulation of the dopamine terminals in the NAc core during training, resulted in impaired conditioned inhibition learning in the ChR2 group (RM ANOVA main effect of cue type F_(1, 14)_= 0.69 p=0.4188, n=20). **(i)** The NpHR group showed robust learning during training following the inhibition of the NAc core dopamine response at the time of omitted shock following the Fear+ Omission cue presentations. (RM ANOVA main effect of cue type F_(1, 10)_= 7.58 p=0.0042, n=20). Data represented as mean ± S.E.M. * *p* < 0.05, ** *p* < 0.01; *** *p* < 0.001; **** *p* < 0.0001; ns = not significant.

First, we recorded dopamine responses to the Fear cue and the Fear+Omission cue following training (**Figure 1g**). The dopamine response to the Fear cue was not different than the response to the Fear+Omission cue (**Figure 1h-i**), even though the predictions and the behavioral responses to these cues differed. This suggests that NAc core dopamine responses do not simply represent the associative strength of the cue or the negative valence of the Fear cue as some have previously hypothesized(28).

Next, we aimed to define how the dopamine response during the shock period (or in response to the shock omission) may play a more critical role in how animals behave and learn across trial types. Previously, we found a positive, significant dopamine response at the time of an omitted footshock; however, this response was smaller than the dopamine response on trials where shock was presented(17). Thus, the biggest positive dopamine response was when the aversive stimulus itself was present, and therefore the dopamine signal could not be explained as a reward-based signal that occurs during omission to signal safety/relief. However, the signal that occurs at the time of omission is also likely an important aspect of dopamine signaling during this task. To this end, we defined how dopamine responses during the shock period changed based on predictions and how this related to future learned behavior in a causal fashion. After initial training, animals were run through a subsequent test session where each cue type was presented; however, no foot shocks were presented. We recorded dopamine release at the time of the predicted, but absent, footshocks on each trial type (**Figure 1j**).

First, confirming our earlier results(17), we showed that there was a positive dopamine response that occurred at the offset of the Fear cue - when the expected footshock would have been presented but was omitted (**Figure 1k**). Interestingly, the size of this response was larger following the Fear cue alone as compared to the Fear+Omission cue (**Figure 1l**). Thus, dopamine release during an omission was highly influenced by the prediction. When the footshock was predicted more strongly by a cue – i.e. following the Fear cue – dopamine responses were larger (**Figure 1k-l**; see **Figure S3** for averaged dopamine responses for individual animals).

Together, these data suggest that rather than a positive valence signal, dopamine transmits a signal that denotes important events to allow for adaptive learning and updating of future behavior. Critically, this includes the omission of predicted aversive events. Overall, these results show that 1) NAc dopamine responds to expected but absent aversive outcomes and 2) this signal is influenced by how strongly the threat is perceived or anticipated.

### Dopamine release in response to predicted aversive events – even in their absence - is necessary and sufficient for learning

Our initial results provide strong evidence suggesting that NAc core dopamine is involved in aversive learning in a valence-free fashion, especially as it relates to predictions about when aversive stimuli will and will not occur. This is critically important, as the ability to predict threats and safety in the environment is an evolutionarily conserved adaptive behavior that is dysregulated by a range of psychiatric disease states. We next wanted to answer two questions: 1) is this dopamine signal in response to an omitted footshock causal to learning and 2) are these effects explained by dopamine as a reward-based signal or a more general perceived saliency signal?

Based on the data in **Figure 1**, there are two possible explanations for what type of signal is being transmitted within these dopamine release signatures. The first possibility is that dopamine release signals errors in reward-based predictions – this is termed reward prediction error [RPE(29,30)]. The other is that dopamine is involved in transmitting a saliency signal that influences how important events are perceived based on the intensity of stimuli and novelty in the environment – termed perceived saliency(17,31). In the RPE account, the positive dopamine signal at the time of omitted shock is a reward-based signal that notes that the outcome is better than expected. In the perceived saliency account, the signal is a valence-free signal that scales with intensity (and predicted intensity) of a stimulus to note that it is an important and unexpected event regardless of valence. Differentiating these hypotheses is particularly critical to understanding the role of dopamine in aversive learning as these accounts would hypothesize opposite effects of dopamine on future behavior (described in detail below).

To parse which of these hypotheses can best explain the data, we enhanced dopamine release using photostimulation of dopamine terminals in the NAc core at the time of the omitted footshock following only the Fear+Omission cue trials during training (**Figure 2a-c; Figure S4**). If NAc core dopamine signals RPE, we should observe a greater difference in freezing behavior between the Fear cue trials and the Fear+Omission trials (i.e., enhanced conditioned inhibition learning) – as this would enhance the positive, reward-based error that occurs when the animals learn that the shock that is expected is not presented (a positive experience). However, if dopamine release transmits a perceived saliency signal, this should attenuate the reduced freezing that develops over time to the Fear+Omission cue (i.e., impaired conditioned inhibition learning). This is because we would be preventing the ability of the signal to diminish at the time of the predicted but absent shock following the cue that signals omission. This reduction in dopamine response is important for animals to learn that the omission is less salient because it was predicted, and a behavioral response is not necessary on future trials of this type.

Confirming our hypotheses of dopaminergic signaling of perceived saliency, our results demonstrated that the group that received optogenetic stimulation of dopamine release (ChR2) at the time of omitted shock following the Fear+Omission cue showed enhanced freezing to the Fear+Omission cue. Because of this, the expected difference between the Fear cue and Fear+Omission cue was not present (**Figure 2d-f, h**) as compared to the eYFP controls (**Figure 2d-f, g**). This effect was persistent during the subsequent testing session (even in the absence of any further optogenetic stimulation; **Figure 2f**) demonstrating that this signal is critical to future learning of the conditioned association, rather than an effect that occurs only at the time of stimulation on those trials (**Figure 2h**).

The converse was also true. In mice that received optogenetic photoinhibition of NAc core dopamine release specifically at the time of the expected but absent footshock (NpHR), there was enhanced learning – i.e. a larger decrease in freezing to the Fear+Omission cue (**Figure 2d-f, i**). This is opposite to the impaired learning that would be expected if the signal was a safety or reward-based signal. These results demonstrate that dopamine responses that occur during the omitted shock causally determine future conditioned behavioral responses in a way that can only be predicted by perceived saliency accounts.

### Dopamine responses during omitted aversive events alter future dopamine release and conditioned behavior

Our results demonstrate that we can control aversive learning by altering the dopamine response elicited by an expected but absent footshock during training. However, if this acts to alter the trajectory of aversive associations that develop over learning, these manipulations should also alter dopamine responses at the time of the omitted but expected shock in the future. To test whether the behavioral results we obtained with optogenetics are due to learning and result in long-term alterations of the dopamine response at the time of the expected, but absent footshock, we combined *in vivo* fiber photometry with optogenetic manipulations in the same animals. Using this approach, we manipulated dopamine terminals in the NAc core at the time of the omitted shock during initial learning and recorded dopamine responses in the same animal in subsequent behavioral test sessions.

Employing the same optogenetic manipulation strategy as above (but this time via the red-shifted excitatory opsin Chrimson), we stimulated NAc core dopamine terminals specifically at the offset of the Fear+Omission cue during training – at the time of the omitted shock (**Figure 3a**; **Figure S5-6**). Next, we recorded dopamine responses in a subsequent test session to both trials in which the Fear cue and the Fear+Omission cue were presented. In test sessions, no shocks were presented. First, we replicated the original results showing that optogenetic stimulation of dopamine terminals at the time of the omitted footshock impaired learning during training (**Figure 3b**; **Figure S7**) and eliminated the differential freezing responses between the Fear cue and Fear+Omission cue (**Figure 3c**). Critically, we also showed that the difference between the dopamine response to the omitted shock following the Fear cue versus following the Fear+Omission cue we demonstrated in **Figure 1** was no longer present after the optogenetic stimulation (**Figure 3d-e**; see **Figure S8** for averaged dopamine responses for individual animals). Thus, here we show that the dopamine response to the expected footshock fundamentally determines both current and future response by scaling with the intensity of the predicted aversive outcome, even in the absence of the actual stimulus.

**Figure 3.**
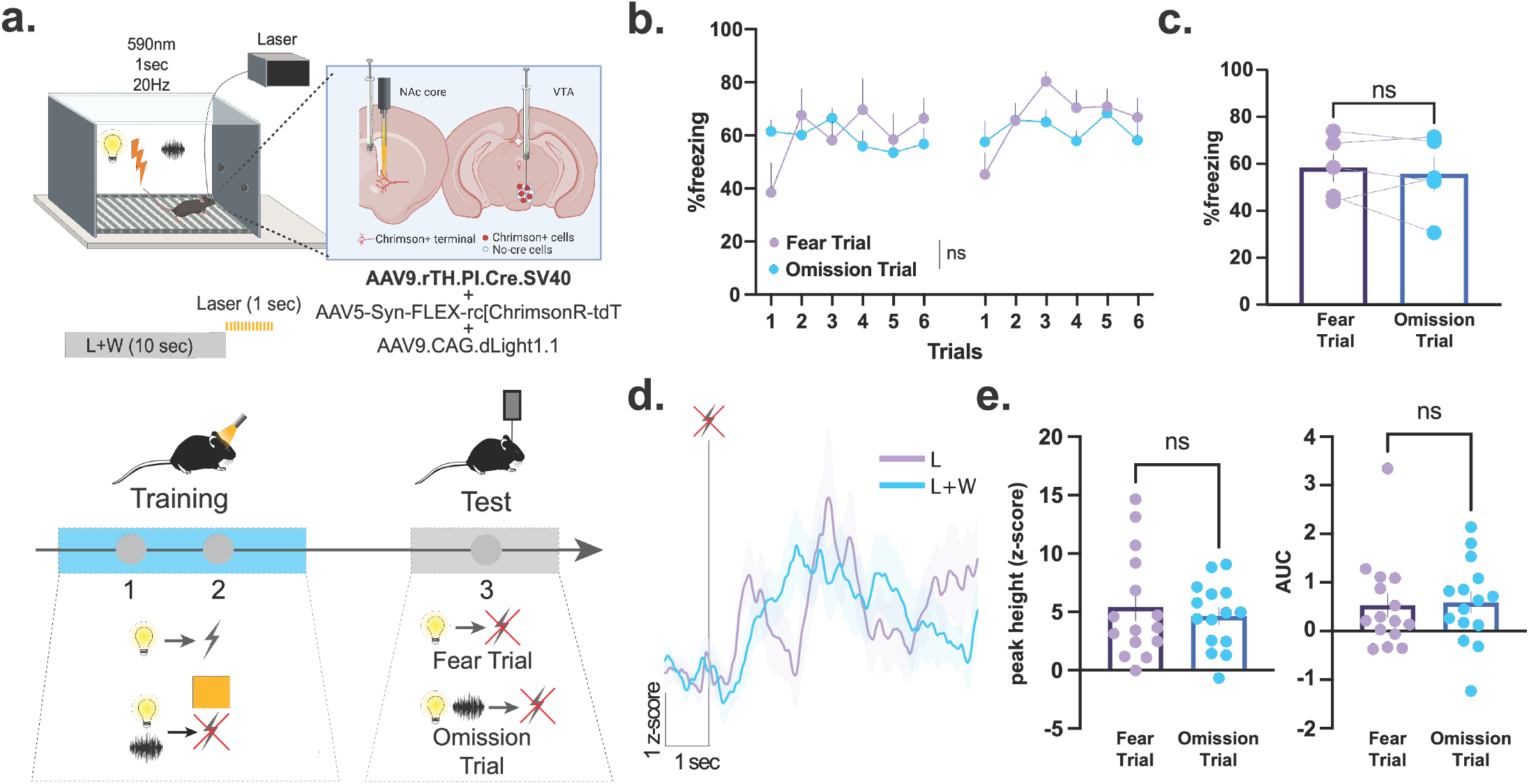
Dopamine release to footshock omission alters future dopamine release and conditioned behavioral responses. **(a)** In mice (n=5), AAV5-Syn-FLEX-rc[ChrimsonR-tdT] was co-injected with AAV9.rTH.PI.Cre into the VTA to achieve dopamine-specific expression of excitatory opsins. AAV9.CAG.dLight1.1 was injected in the NAc core for concurrent dopamine imaging in the same animals. Dopamine terminals were optogenetically photostimulated (via Chrimson) at the time of the omitted shock following the Fear+Omission cue (houselight+whitenoise). Photostimulation of dopamine terminals during training disrupted conditioned inhibition learning. **(b)** Freezing response to the Fear cue only did not differ from the freezing response to the Fear+Omission cue across the training sessions (RM ANOVA main effect of group F_(1, 7)_= 1.13 p=0.3232, n=5) and **(c)** during the subsequent test session (paired t-test, *t*_4_=0.62, *p*=0.5684, n=5 mice). **(d)** Averaged dopamine responses to the Fear cue versus Fear+Omission cue. **(e)** Peak height (Nested t-test, *t*_28_=0.56, *p*=0.5789, n=5 mice) and area under the curve (AUC; Nested t-test, *t*_8_=1.14, *p*=0.8878, n=5 mice) of the dopamine responses. Data represented as mean ± S.E.M., ns = not significant.

### The effects of dopamine terminal photostimulation can be explained by perceived saliency, but not reward prediction, or the associative value of cues

The idea of dopamine in the NAc core as a perceived saliency signal stems from the theoretical framework we published recently, which is computationally represented as a formal model of associative learning (the Kutlu-Calipari-Schmajuk [KCS] model(17); **Figure 4a**). The KCS model describes theoretically how we allocate attention in an environment. Importantly, this is heavily influenced by both the physical intensity (how physically strong a stimulus is) and novelty in an environment. In this way, things that are both physically intense (like a strong footshock) and things that are novel (like a prediction not being met) combine to elicit a response that scales with how much attention needs to be allocated to that event or stimulus in order for adaptive behavior to occur. This means that dopamine can be elicited by both stimuli that are present and by stimuli that are predicted but absent – which is the signal that we observed here is causally related to behavior. Thus, the results of the dopamine recordings and optogenetic manipulation experiments presented here suggest that the dopamine response we observe aligns closely with perceived saliency; however, we wanted to empirically test this hypothesis.

**Figure 4.**
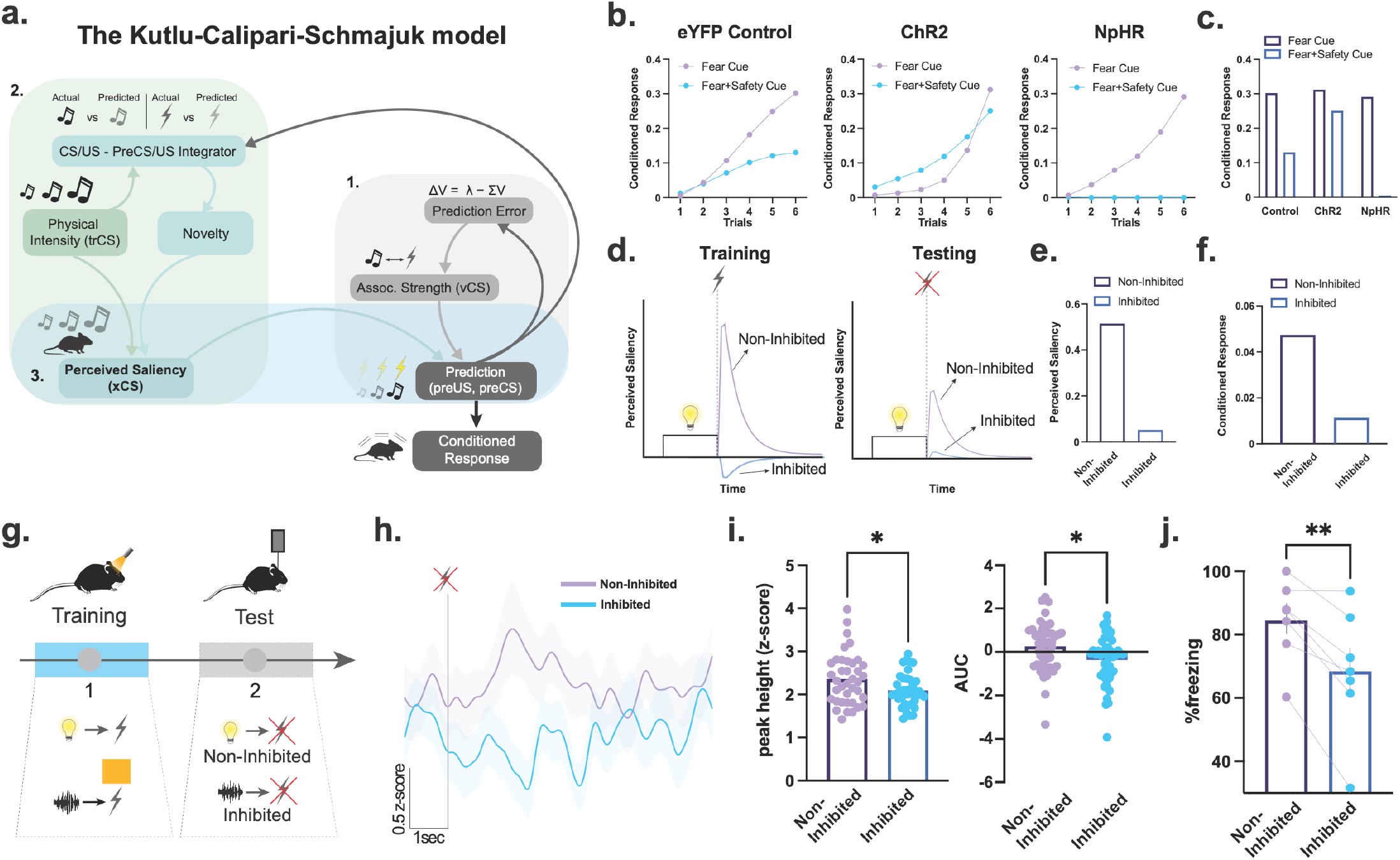
NAc dopamine responses and associated behavior can only be explained by dopamine transmitting a perceived saliency signal. **(a)** The Kutlu-Calipari-Schmajuk (KCS) model. The model has 4 core components. *1) Associative component:* Based on a Rescorla-Wager type prediction error term. *2) Attentional component:* Mismatch between predicted/unpredicted stimuli increases novelty, and in turn, attention to all stimuli in the environment. 3) *Perceived Saliency:* Novelty, attention, and the physical intensity of a stimulus determine perceived saliency. 4) *Behavioral response component:* Perceived saliency is combined with associative strength to produce a prediction of an outcome. **(b)** Simulations of the KCS model during training and **(c)** testing of conditioned inhibition. Our simulations show that the KCS model accurately mimics conditioned inhibition (i.e. weaker conditioned response to the Fear cue compared to the Fear+Omission cue). When the perceived saliency of the outcome (footshock) was set to a higher value to mimic the optogenetic stimulation of NAc dopamine (ChR2), similar to our experimental data, the KCS model fails to learn conditioned inhibition as we observed in **Figure 2&3**. Conversely, when the perceived saliency was set to a negative value to mimic optogenetic inhibition of dopamine terminals, the KCS model exhibits more robust safety learning as we observed in **Figure 2. (d)** KCS model simulations where the perceived saliency of the footshock is set to a negative value during a cue-outcome training to mimic optogenetic inhibition of NAc core dopamine. **(e)** Model simulations demonstrate that following artificial inhibition of perceived saliency during training, perceived saliency of the predicted but absent outcome becomes weaker. **(f)** The model predicts that following optogenetic inhibition of the dopamine response to the outcome the conditioned response to the inhibited cue will be weaker compared to a non-inhibited cue. **(g)** In order to empirically test KCS model’s predictions, mice (n=7) were trained where two separate cues each predicted the presentation of a footshock. Dopamine release during the footshock was optogenetically inhibited following one cue (Inhibited Cue), but not the other (Non-Inhibited Cue). **(h)** Dopamine response at the time of the omitted shock following the inhibited cue (during the expected but absent footshock) was weaker compared to the signal following the Non-inhibited cue. **(i)** The peak-height (Nested t-test, *t*_78_=2.13, *p*=0.0357, n=7 mice) and the area under the curve (AUC; Nested t-test, *t*_78_=2.43, *p*=0.0173, n=7 mice) of the dopamine response to the expected but absent footshock at the offset of the Inhibited cue was smaller compared to the Non-inhibited cue. **(j)** As predicted by the KCS model simulations, the freezing response to the Non-inhibited cue was stronger than the Inhibited cue response (paired t-test, *t*_6_=4.42, *p*=0.0044, n=7 mice). Data represented as mean ± S.E.M. * *p* < 0.05, ** *p* < 0.01.

One of the advantages of the KCS model is that it allows us to simulate different behavioral paradigms and model the trajectory of learning as animals form aversive associations. Here we simulated the conditioned inhibition paradigm we employed above using the KCS model. Then, we computationally manipulated the perceived saliency and prediction error terms in the model in order to determine if we would get the same behavioral effect as we saw when we optogenetically increased or decreased dopamine (a hypothesized perceived saliency signal).

The model was able to mimic the behavioral responses that occur in this paradigm in control animals (**Figure 4b**; eYFP controls). Moreover, by assuming that the optogenetic stimulation and inhibition of the NAc core dopamine terminals increases and decreases the perceived saliency of the expected but absent footshock, respectively, the model was also able to capture the specific effects of these optogenetic manipulations. That is, the model predicted impaired conditioned inhibition learning when the perceived saliency was increased (**Figure 4b**; ChR2) and also predicted enhanced conditioned inhibition learning as a result of inhibited perceived saliency of the expected but absent footshock (**Figure 4b**; NpHR). When we computationally manipulated the prediction error for the KCS conditioned inhibition simulations we observed the opposite effect on behavior, where artificially increasing the prediction error resulted in faster conditioned inhibition learning whereas the model failed to learn conditioned inhibition when the prediction error was suppressed (**Figure S9**). This was inconsistent with the experimental data we observed throughout the manuscript. These simulations give theoretical support for the experimental results we obtained here and places them well within our framework of dopaminergic encoding of perceived saliency.

### Optogenetically inhibiting dopamine at the time of the shock prevents the formation of aversive associations

The KCS model simulations also offer testable predictions regarding the source for the signaling of perceived saliency of expected outcomes by dopamine. Simulations showed that when the perceived saliency of the footshock is computationally inhibited, the subsequent perceived saliency of the expected footshock is diminished **(Figure 4d-e**; **Figure S10**) and therefore, the conditioned response to that cue will also be weaker compared to a “non-inhibited” cue (**Figure 4f**). Our simulations also demonstrated that this is due to the fact that with diminished perceived saliency of the footshock during training, the associative strength between the inhibited cue and the outcome is weaker than the strength of the association between the non-inhibited cue and the outcome (**Figure S11-12**), thus the conditioned behavioral response should also be reduced.

Therefore, according to the KCS model, if the perceived saliency of the footshock is suppressed during an initial cue-footshock pairing, the association between those two stimuli will not be formed adequately. In this framework, positive dopamine responses that occurs at the time of the shock is necessary for the aversive association. This is in opposition to a reward-based idea of dopamine coding which would suggest that the dopamine responses to an aversive footshock should be negative to signal the negative valence of the shock. Thus, decreasing dopamine in response to a shock would increase the rate of learning if dopamine transmitted a reward or valence-based signal. We empirically tested this.

We paired two different cues with a footshock during training (**Figure 4g**). We optogenetically inhibited NAc core release during the footshock presentation following one cue (inhibited cue) but not the other (non-inhibited cue). Then, we recorded dopamine responses at the time of the expected but absent footshock during a subsequent test session (**Figure 4g; Figure S13**). We found that following the inhibited cue presentation, the dopamine response to the expected but absent footshock was weaker compared to the non-inhibited cue (**Figure 4h-i**; see **Figure S14** for averaged dopamine responses for individual animals). Furthermore, as predicted by the model (**Figure 4f**), the freezing response to the inhibited cue was also weaker than the freezing response to the non-inhibited cue (**Figure 4j**). Thus, positive dopamine responses that occur at the time of the shock are necessary for the development of an aversive conditioned behavioral response in a way that can only be explained by dopamine as a perceived saliency signal.

Moreover, supporting our hypothesis that NAc core dopamine release does not directly represent prediction/expectation of outcomes at the level of cue encoding, we found that the dopamine response during the inhibited and non-inhibited cues did not differ (**Figure S15**). These results suggest that the dopaminergic encoding of the perceived saliency of aversive outcomes are required to form associations between cues and outcomes and the resulting conditioned responses; although dopamine is not driving the associative value that is acquired by the cue itself.

Together, our results show that dopamine responds to predicted, but absent, aversive stimuli. This response scales with the strength of their prediction. That is, the stronger the prediction that that aversive outcome will occur, the stronger the dopamine response will be if it is omitted. However, this is not represented in the dopamine response to the cue itself, which did not similarly scale to the same level with these predictions. This is particularly important as it helps to explain how dopamine is specifically involved in predicting when threats will and will not occur in an environment– something that is fundamental to survival and is dysregulated in many anxiety and stress disorders. These data are also important as they diverge from the traditional theoretical understanding of dopaminergic information encoding as a reward-based prediction signal and show that the effects of dopamine on aversive learning are better explained by dopamine release as a perceived saliency signal.

## Discussion

Here we show that dopamine release in the NAc core plays a causal role in guiding animals to learn to predict when aversive stimuli will and will not be presented in an environment. Critically, we show that this occurs through the modulation of dopamine release at the time of the predicted outcome (shock or omission). Interestingly, the difference in these predictions was not robustly represented in the dopamine response to the cue itself. Further, the dopamine response that occurred at the time of an omitted footshock scaled with the prediction of that outcome – with larger dopamine responses occurring when the omitted shock was more strongly predicted by the antecedent cue. Thus, we find that dopamine release is modulated by predictions, but does not seem to track the prediction itself. Finally, we show that dopamine release signatures can be explained by dopamine transmitting a perceived saliency signal, but not an associative or reward-based signal. This is particularly important as it helps to explain how dopamine is specifically involved in predicting when threats will and will not occur in an environment – a process that is fundamental to survival and is dysregulated in many anxiety and stress disorders(3–5). Overall, this study elucidates the role of NAc core dopamine release in aversive learning in a theory-based and stimulus-specific fashion and offer potential avenues for understanding the neural mechanisms involved in anxiety and stress disorders where this behavioral process is dysregulated.

We are not the first to show that dopamine plays a critical role in aversive learning; however, some of the previous work in this area has still concluded that dopamine release in the NAc core is involved the processing of relief, or safety, and thus could still be due to positive valence coding. For example, studies from our lab and others showed that avoidance of aversive stimuli during active avoidance learning (or safety learning) is associated with a positive dopamine response in the NAc(10,11,17,20). Further, dopamine is evoked by the omission of a predicted aversive stimulus in Pavlovian fear conditioning tasks(17). To this end, some have hypothesized that the avoidance or omission of an aversive stimulus is a relief/safety/reward-based signal that guides future behavior(10,11,32). However, more recent work has shown that the NAc core is causally linked to aversive learning in a way that cannot be predicted by reward or valence processing. For example, dopamine is released in response to both appetitive and aversive stimuli and scales positively with intensity in both cases(17,33). Further, dopamine release in the NAc core to cues that predict the presentation of an aversive stimulus after an operant response (punishment) are also positive, rather than negative to signal negative valence(17). In addition, our results presented here also refute the “relief hypothesis” of dopamine(34). Specifically, we found that optogenetically increasing this signal during the omission of the expected aversive outcome impaired the ability of animals to learn about when aversive stimuli would not be presented (similar to safety learning paradigms), rather than facilitating it as one would expect as a result of a larger relief from threat.

Similarly, our results offer additional evidence against the widely discussed RPE hypothesis of dopamine in learning in memory. Previous literature suggests that dopamine cell bodies within the VTA signal a prediction error specific to reward outcomes(6,7,30,35). Since these neurons are the source of dopamine release in projection targets such as the NAc core, the assumption has been that dopamine release also obeys the same coding rules and, therefore, signals RPE. However, there is now evidence convincingly demonstrating that the role of dopamine released in the NAc goes beyond simply signaling RPE as it is heavily involved in signaling aversive stimuli and outcomes(11,17–19). One important data point from this study is the observation that dopamine responds at the time of expected stimulus, even when the expected stimulus is now omitted. This provides testable predictions to compare the RPE hypotheses to more contemporary models that have suggested that dopamine release in the NAc functions as a perceived saliency signal(17,22,24). If dopamine represents any kind of prediction error (reward-specific or valence-free), enhancing this signal should also enhance the learning rate of cue-outcome associations(36). However, we found the opposite effect; optogenetically stimulating the dopamine response to the omitted footshock following the Fear+Omission cue did not enhance the reduced freezing that should occur in response to the cue that signals shock omission. In fact, stimulation of dopamine release resulted in impaired inhibitory learning. This is consistent with our previous results showing that enhancing the dopamine response to the expected but absent footshock impairs fear extinction(17). Thus, dopamine release causally mediates associative learning; however, the outcomes of these experiments are best explained by dopamine as a perceived saliency signal.

One particularly important aspect of these data is the fact that dopamine release was modulated by trial type at the time of the omitted footshock, but not to the predictive cue. For example, dopamine release to an omitted footshock was largest when the antecedent cue predicted that a shock would occur more strongly, and smallest when a cue that predicted the shock would not occur was presented. However, the response to the cue itself between these two trial types did not differ. Therefore, while the effects of dopamine in these tasks are influenced by predictions, the dopamine signal does not itself encode the aversive prediction at the level of the cue response. This gives further support to the idea that dopamine release in the NAc core transmits a perceived saliency signal, rather than a reward-based prediction signal. Importantly, the perceived saliency of stimuli and events is heavily influenced by the prediction provided by the antecedent predictive cue, which can alter the attention an animal allocates to that stimulus in an environment (31,37). This effect explains why these dopamine signals are modulated by predictions, and in many cases look like they encode the prediction itself(29,38,40,42), when in fact they are dissociable from one another under key conditions. Together these findings put an emphasis on the importance of understanding the nuanced relationship between observed neural signals and behavioral responses.

This work is also particularly important in the context of stress disorders. Anxiety and stress disorders, including but not limited to panic disorders, phobias, generalized anxiety disorder, and post-traumatic stress disorder (PTSD), are among the most prevalent mental disorders, affecting 40 million Americans with an 18% 12-month prevalence and a 30% lifetime prevalence(39,41). One of the major symptoms of anxiety and stress disorders such as post-traumatic stress disorder (PTSD) is negative emotional responses such as re-experiencing the traumatic events, avoidance of trauma-related memories and hyperarousal, which persist in novel contexts that were previously perceived as safe(43,44). Studies suggest a strong link between the ability to learn that cues previously paired with danger no longer signal threat, known as extinction learning, with anxiety and stress disorders as this form of learning is delayed in individuals with anxiety disorders(3–5). Furthermore, patients with anxiety disorders also show impaired safety discrimination, a form of inhibitory learning where a stimulus or a context signals safety versus danger, and impaired ability to transfer learned inhibition to a fear-eliciting stimulus(2,45–48). Therefore, it is largely accepted that deficits in fear response inhibition may be one of the main attributes of anxiety and stress disorder symptomatology(49,50). In support of a role of impaired safety learning in anxiety disorders, several studies found that war veterans(47) and the civilian population(46) with PTSD fail to learn associations between stimuli and safety. Moreover, there is evidence showing that children that exhibit deficits in learning safety signals are at a higher risk of developing anxiety disorders when they become adults(51). These studies suggest a causal relationship between impaired safety learning and anxiety and stress disorders and, therefore, our results suggesting that dopamine response in the NAc core causally determines safety learning have important implications for human psychopathologies and potential treatment options for those conditions.

Together, we show that NAc core dopamine responses to expected but omitted aversive stimuli causally determine associative learning for aversive stimuli in mice. These results add to the growing literature supporting the dopaminergic encoding of perceived saliency by dopamine in the NAc. Our conclusions regarding the dopaminergic information encoding in associative learning also have important clinical implications for anxiety and stress disorders. As mentioned earlier, impaired safety learning is one of the major symptoms of anxiety and stress disorders such as PTSD(2,43–48,51), and the involvement of the dopaminergic processes has been implicated in these disorders(13–15,52). Together with a recent human study(53), our results suggest that temporally specific suppression of dopamine during the omission of aversive stimuli (or in safety learning contexts) may have beneficial effects in clinical populations. In sum, these results have far-reaching implications for the theory of learning and memory, the understanding of the mesolimbic neurocircuitry, and the psychopathology of anxiety and stress disorders.

## Acknowledgements

This work was supported by NIH grants KL2TR002245 to M.G.K, DA042111 and DA048931 to E.S.C., GM07628 to J.E.Z. as well as by funds from the VUMC Faculty Research Scholar Award to M.G.K., the Pfeil Foundation to M.G.K., Brain and Behavior Research Foundation to M.G.K, and E.S.C, the Whitehall Foundation to E.S.C., and the Edward Mallinckrodt, Jr. Foundation to E.S.C.

## Supplementary Methods

### Apparatus

For all mouse fiber photometry and optogenetic experiments, animals were trained and tested daily in individual operant conditioning chambers (Med Associates Inc., St. Albans, Vermont) fitted with visual and auditory stimuli including a standard house light, a white noise generator, and a 16-tone generator capable of outputting frequencies between 1 and 20 KHz (85 dB).

### Histology

Mice were deeply anaesthetized with an intraperitoneal injection of a ketamine/xylazine mixture (100 mg/kg;10 mg/kg) before being transcardially perfused with 10 mL of 1x PBS solution followed by 10 mL of cold 4% PFA in 1x PBS. Animals were subsequently decapitated, and the brain was extracted and postfixed in the 4% PFA solution stored at 4 °C for at least 48 hours before being dehydrated in a 30% sucrose in 1x PBS solution stored at 4 °C. After sinking, tissue was sectioned (35 μm slices) on a freezing sliding microtome (Leica SM2010R) and then placed in a cryoprotectant solution (7.5% sucrose + 15% ethylene glycol in 0.1 M PB) stored at −20 °C until immunohistochemical processing. For the optogenetic experiments using AAV9.rTH.PI.Cre.SV40, we also validated the targeting of TH+ cells in the VTA via an anti-TH antibody (mouse anti-TH; Millipore #MAB318, 1:100). Sections were then incubated with secondary antibodies [gfp: goat anti-chicken AlexaFluor 488 (Life Technologies #A-11039), 1:1000 and TH: donkey anti-mouse AlexaFluor 594 (Life Technologies # A-21203), 1:1000] for 2 h at room temperature. After washing, sections were incubated for 5 min with DAPI (NucBlue, Invitrogen) to achieve counterstaining of nuclei before mounting in Prolong Gold (Invitrogen). Sections were mounted on glass microscope slides with ProLong Gold antifade reagent. Fluorescent imaging was conducted using a BZ-X700 inverted fluorescence microscope (Keyence) under a dry 20x objective (Nikon). Injection site locations and optical fiber placements were determined with serial images in all experimental animals.

### Computational modeling

The Kutlu-Calipari-Schmajuk model (KCS) has been developed based on an attentional neural network model of Pavlovian conditioning (Schmajuk-Lam-Gray-Kutlu model, SLGK model(1,2)). At the core of the basic model (depicted in **Figure 4a**; see below for complete list of equations) is an error prediction term where associations between multiple conditioned stimuli (VCS-CS), as well as between conditioned and unconditioned (VCS-US) stimuli, are formed based on the same Rescorla-Wagner-based predictions. In addition to the associative components, an attentional “Perceived Saliency” term is included to provide a representation of what is being predicted by a conditioned stimulus when the stimulus is physically present as well as when it is absent. The model assumes that Perceived Saliency of a stimulus is determined by stimulus saliency combined with the level of attention directed to a given stimulus. This way, two stimuli with equivalent physical intensity (e.g., two tones with equal dB values) can be weighted differently and receive processing priority when forming associations with an outcome depending on their attentional value(3). Perceived Saliency is computationally defined as the product of stimulus saliency (termed ‘CS’ in the model) and attentional value of a stimulus (zCS). The core factor that controls attentional allocation is the level of Novelty in a given context, which is determined by the level of mismatch between predictions and actual occurrences of events on a global scale. Accordingly, the perceived saliency of a stimulus increases when Novelty is high and the organism directs more attention to that stimulus, even when the saliency is constant. One of the most important tenets of the model is that even stimuli that are predicted but absent activate a representation, albeit weaker than the perceived saliency of stimuli that are present. The concept of novelty-driven perceived saliency allows our model to be able to describe learning phenomena where stimuli form associations with other stimuli in their absence (e.g., sensory preconditioning(4)).

The constant values are determining rates of each term described below and are taken from the SLGK model (K1=0.2, K2=0.1, K3=0.005, K4=0.02, K5=0.005, K6=1, K7=2, K8=0.4, K9=0.995, K10=0.995, K11=0.75, K12=0.15, K13=4).

#### Stimulus trace and value

In the KCS model, time is represented as the units (t.u.) wherein stimuli are presented, making time-specific predictions of each component of the model possible. In addition to the duration of active presentation of a stimulus, the model assumes a short-term memory trace represented in time for each stimulus (τCS(5,6)). The memory trace decays after the offset of the stimulus presentation. The strength of the memory trace of an individual stimulus (inCS) is determined by the conditioned stimulus saliency (CS) and strength of the prediction of the CS by other stimuli in the environment (preCS). K1 is the decay rate of the stimulus memory trace:

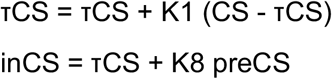

#### Novelty

Novelty of a stimulus (S; CS or US) is proportional to the difference between the actual value (λS and isS) and the prediction of that stimulus (BS and ipreS), also denoted as the CS/US - preCS/US Integrator. Novelty increases when the stimuli are poor predictors of the US, other CSs (i.e., when the US, other CSs, or the context, CX, are underpredicted or overpredicted by the CSs and the CX). Total Novelty (Noveltytotal) is given by the Novelty values of stimuli present in the environment.

NoveltyS ∼ Σ lλS - BSl

Integrator = NoveltyS = lisS - ipreSl isS = K9 isS+Rout (1 - isS)

ipreS = K10 ipreS + ipreS (1 - ipreS)

Noveltytotal = Noveltytotal + NoveltyS

#### Attention

Changes in attention zCS (ΔzCS,-1 > zCS > +1) to an active or predicted CS are proportional to the salience of the CS and are given by:

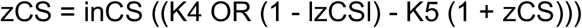

We assume that the orienting response (OR) is a sigmoid function of Novelty

OR = (Noveltytotal^2^/(Noveltytotal^2^ + K11^2^))

ΔzCS > 0; when Novelty > ThresholdCS

ΔzCS < 0; when Novelty < ThresholdCS

ThresholdCS = K5/K4

#### Aggregate Stimulus Prediction

The aggregate prediction of the US by all CSs with representations active at a given time (BUS) is determined by:

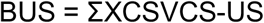

#### Associative Strength

The change in the strength of an association between a CS and a US or between a CS and another CS is determined by:

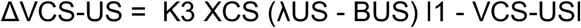

#### Perceived Saliency

The Perceived Saliency of a CS or a US that is present or absent (given by the trace; τCS at a given time point), is proportional to the prediction of (BCS) and attention to that CS (zCS):

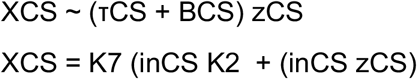

#### Pavlovian Conditioned Response

The US-specific CR is a sigmoid function of the total prediction of the US by all stimuli in the environment (preUS = preUS + BUS):

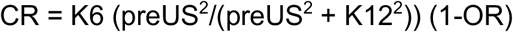

### Behavioral Data Analysis

Statistical analyses were performed using GraphPad Prism (version 8; GraphPad Software, Inc, La Jolla, CA) and Matlab (Mathworks, Natick, MA). Freezing behavior, identified as the time of immobility except respiration during the stimulus duration, was calculated and converted into percent freezing ((freezing time * 100)/ stimulus duration). For all other freezing data, we used a one-way ANOVA followed by Tukey’s post-hoc analysis. All data were depicted as group mean ± standard error of the mean (S.E.M.).

### Fiber Photometry Analysis

The analysis of the fiber photometry data was conducted using a custom Matlab pipeline. Raw 470nm and isosbestic 405nm traces were used to compute ΔF/F values via polynomial curve fitting. For analysis, data was cropped around behavioral events using TTL pulses, and for each experiment, 2s of pre-TTL up to 20 seconds of post-TTL ΔF/F values were analyzed. In order to normalize the signal, we used the isosbestic channel signal (405nm(7)) to calculate our ΔF/F ((ΔF/F =F470-F405)/F405). In addition, all fiber photometry data were converted to and reported as z-scores. We z-scored dopamine signals around the event of interests, such as the CS+, using their own local baseline (2 seconds prior to the cue onset). Z-scores were calculated by taking the pre-TTL ΔF/F values as baseline (z-score = (TTLsignal - b_mean)/b_stdev, where TTL signal is the ΔF/F value for each post-TTL time point, b_mean is the baseline mean, and b_stdev is the baseline standard deviation). This allowed for the determination of dopamine events that occurred at the precise moment of each significant behavioral event. For statistical analysis, we calculated the AUC and peak height. The AUCs were calculated via trapezoidal numerical integration on each of the z-scores across a fixed timescale. The peak height values were the maximum values after the TTL onset. Baseline dopamine responses were calculated as the z-scored dopamine values during the inter-trial interval 20 seconds prior to the CS+ presentations. Unpaired t-tests were employed to test the group differences for all fiber photometry-based dependent variables. We also calculated maximum z-scores for event fiber photometry traces and analyzed them to see if these were significantly different from the critical z-score at p=0.05 level (1.645) using independent-t-tests

---

**Figure S1.**
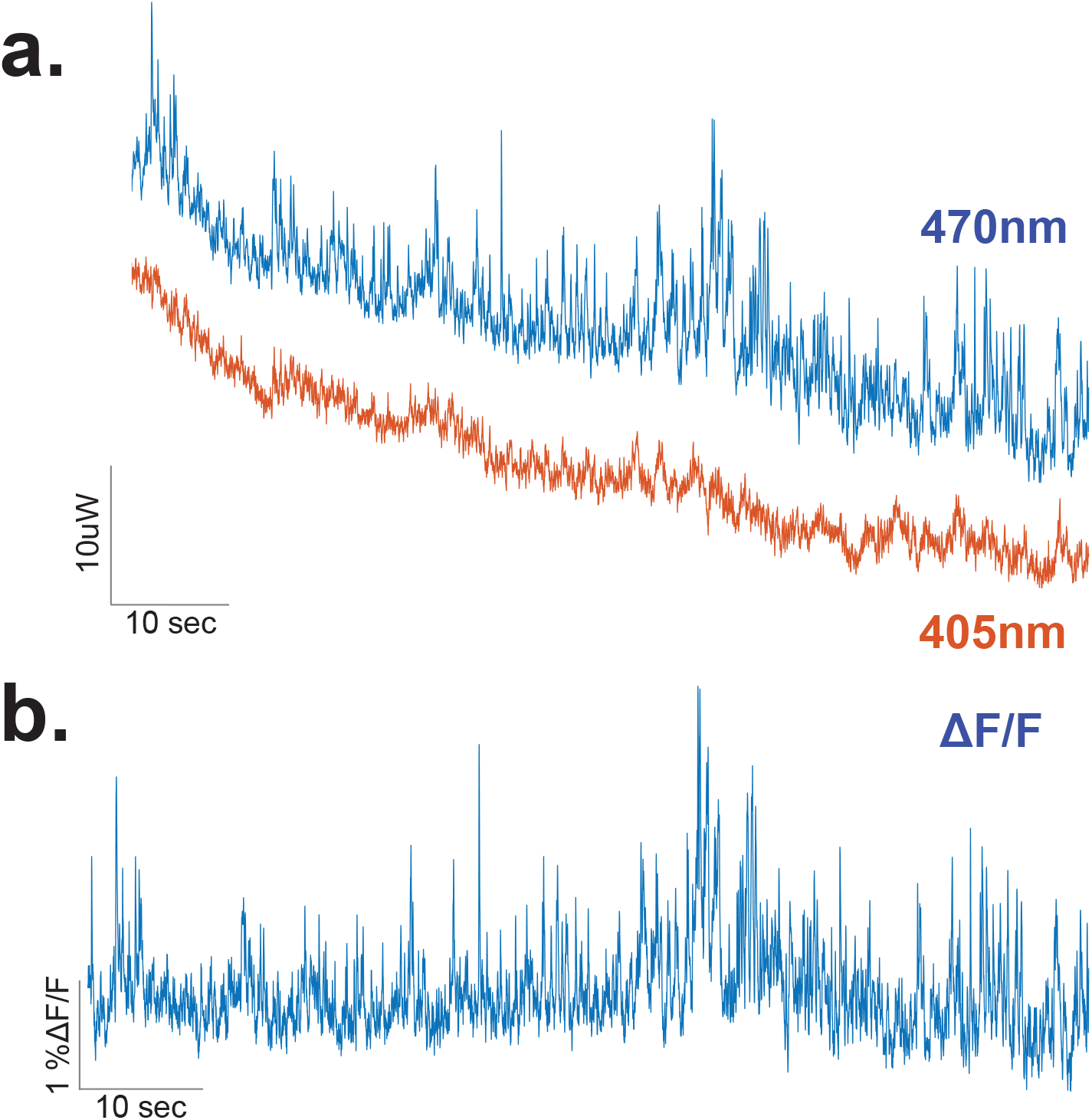
Representative dLight recording traces during training. **(a)** Representative traces for 470 nm excitation (dLight) and 405 nm excitation (isosbestic control) channels in an individual animal at baseline. **(b)** Representative ΔF/F trace showing dopamine transients in the nucleus accumbens core.

**Figure S2.**
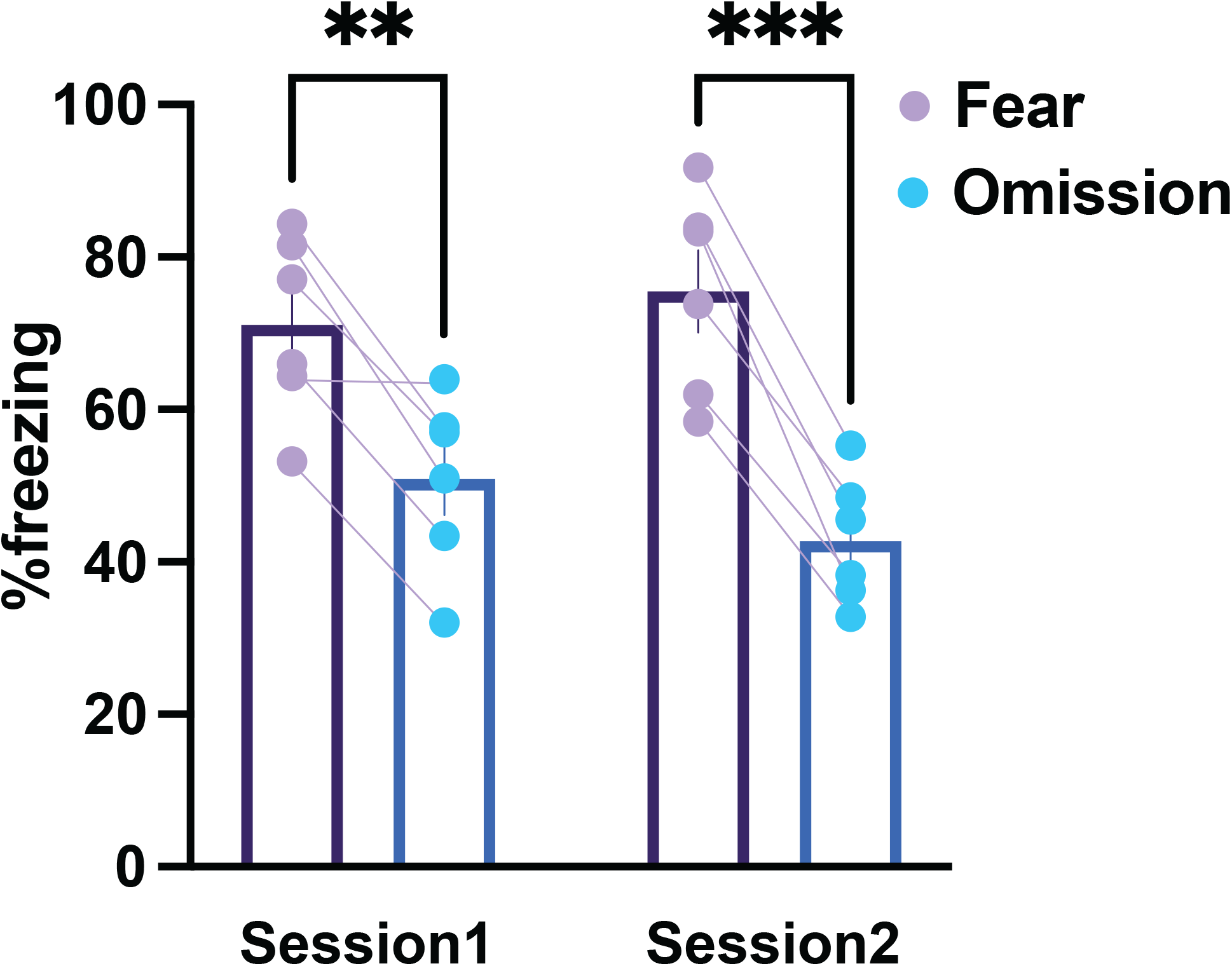
Mice showed robust inhibitory learning during training. Mice showed reduced freezing response to the Fear+Omission cue compared to the Fear cue only trials during both Session 1 (RM ANOVA Cue main effect *F*_(1,5)_= 70.44, *p*=0.0004; Sidak post-hoc p<0.01, n=6 mice) and Session 2 (Sidak post-hoc p<0.001, n=6 mice) of the aversive conditioned inhibition training. Data represented as mean ± S.E.M. ** *p* < 0.01; *** *p* < 0.001.

**Figure S3.**
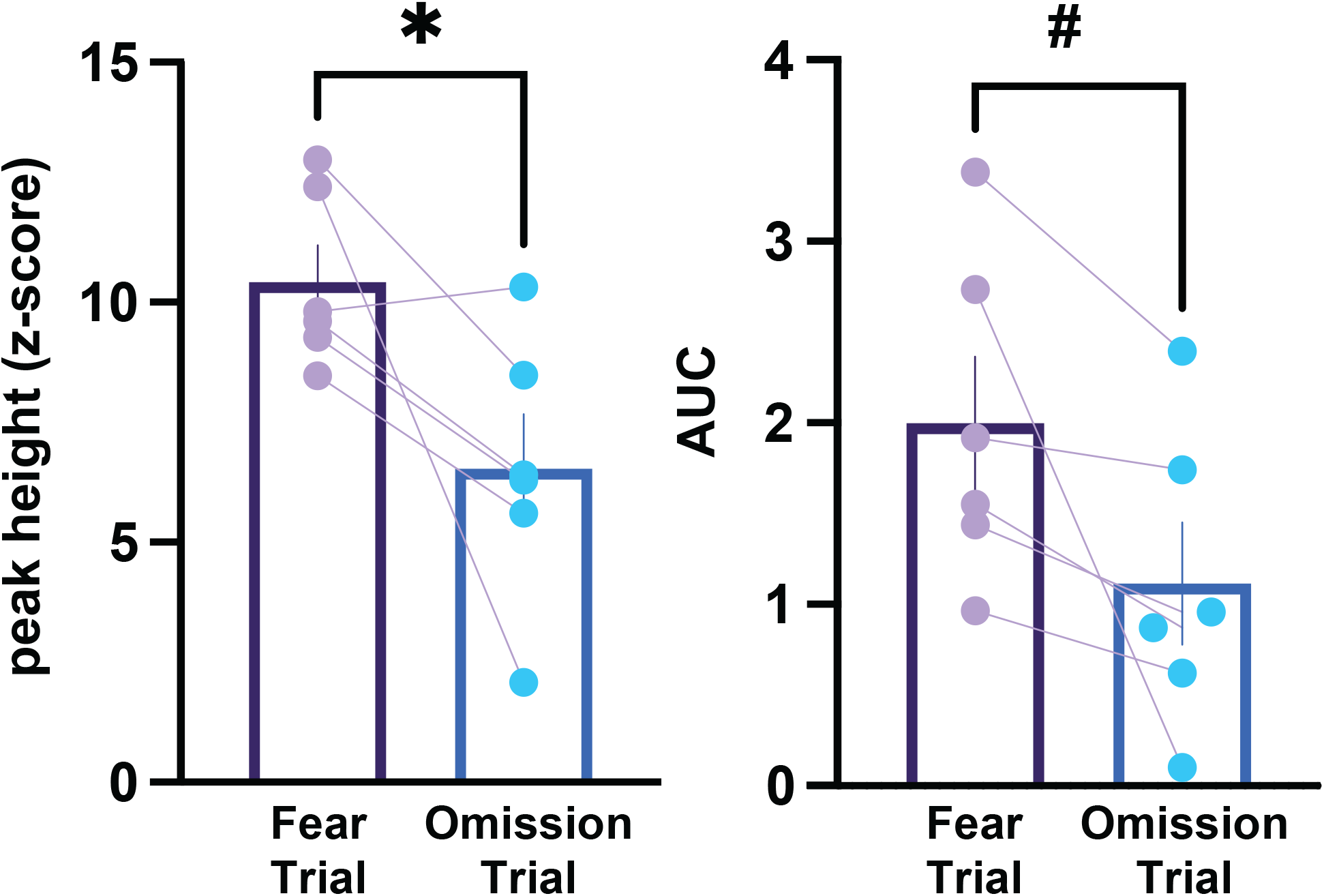
Averaged dopamine responses from individual animals at the offset of the Fear cue – the time of predicted shock – was larger than following the Fear+Omission cue. The peak-height (Paired t-test, *t*_5_=2.67, *p*=0.0442, n=6 mice) and the area under the curve (AUC; Paired t-test, *t*_5_=2.40, *p*=0.0614 n=6 mice) of the dopamine response to the expected but absent footshock at the offset of the Fear cue was larger than the dopamine response at the offset of the Fear+Omission cue. Data represented as mean ± S.E.M. * *p* < 0.05; # *p* = 0.06.

**Figure S4.**
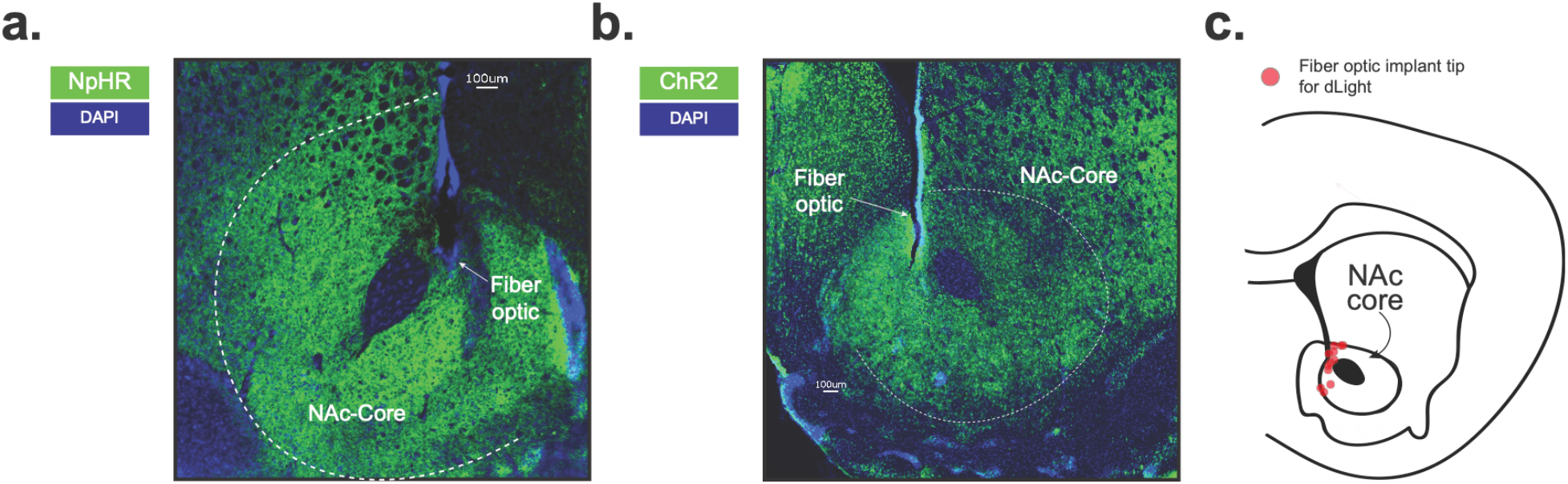
Excitatory and inhibitory opsin expression in dopamine terminals in the NAc core. A fiber optic cannula was placed directly above the NAc core in order to deliver blue or yellow laser stimulation to optogenetically control NAc core dopamine terminals. This was achieved via the expression of an excitatory (AAV5.Ef1a.DIO.hChR2.eYFP; ChR2) **or** an inhibitory (AAV5-Ef1a-DIO.eNpHR.3.0-eYFP; NpHR) opsin expressed in in combination with a TH-specific Cre virus in the VTA (AAV9.rTH.PI.Cre.SV40). **(a)** Representative histology showing viral expression of halorhodopsin (NpHR; green) restricted to the NAc core and **(b)** channelrhodopsin (ChR2; green). **(c)** Schematic showing fiber optic placements in both ChR2 and NpHR animals.

**Figure S5.**
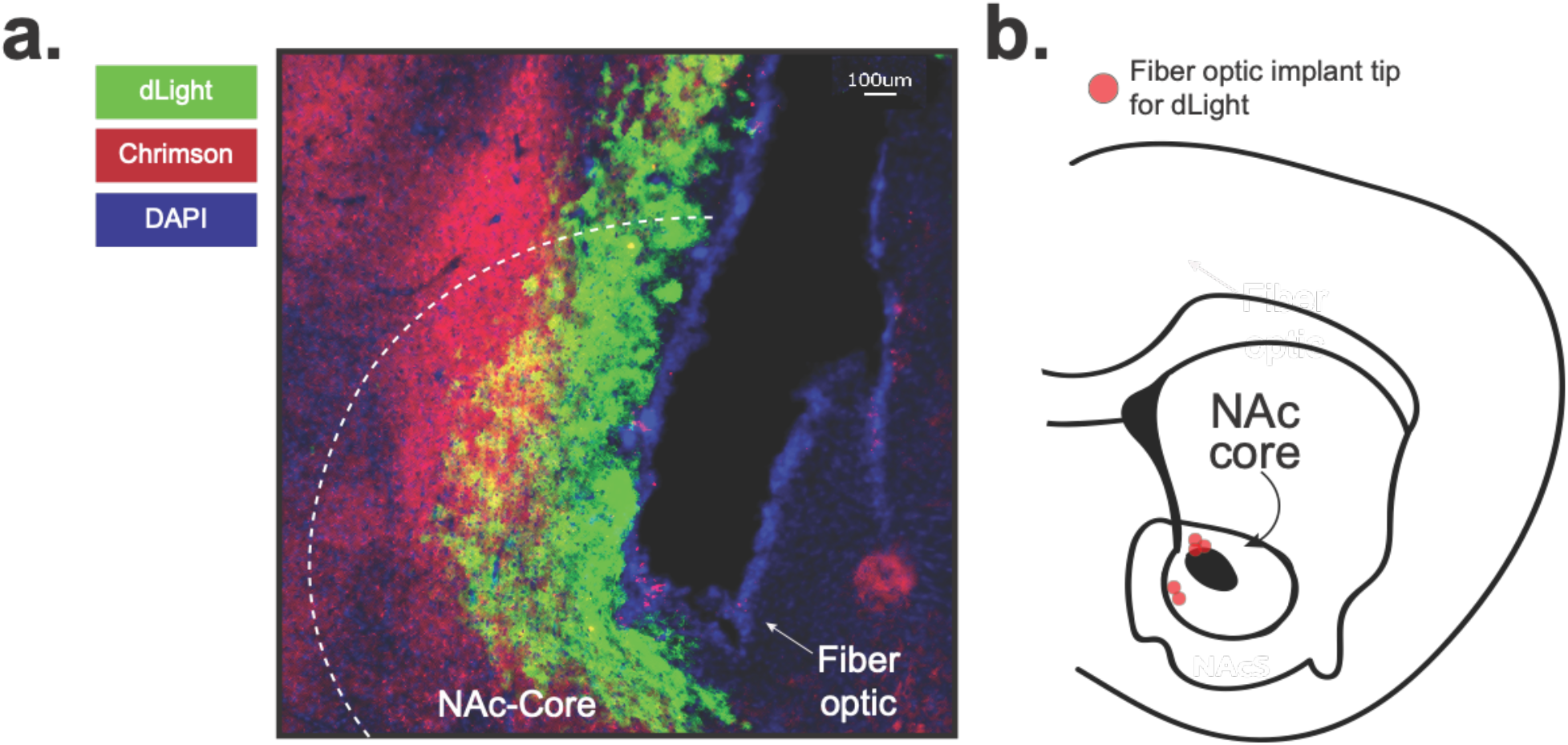
Expression of a red-shifted excitatory opsin (in dopamine terminals) in conjunction with dLight in the NAc core. **(a)** A fiber optic cannula was placed directly above the NAc core in order to record dopamine transients via dLight (AVV5.CAG.dLight1.1). A yellow laser was used to optogenetically control NAc core dopamine terminals via Chrimson. Chrimson expression in dopamine terminals was achieved by expressing (Chrimson.FLEX: AAV5-Syn-FLEX-rc[ChrimsonR-tdTomato]) in combination with a TH-specific Cre expressing virus co-injected into the VTA (AAV9.rTH.PI.Cre.SV40). Representative histology showing viral expression of dLight (green) and Chrimson (red) restricted to the NAc core and **(b)** schematic showing fiber optic placements in Chrimson animals.

**Figure S6.**
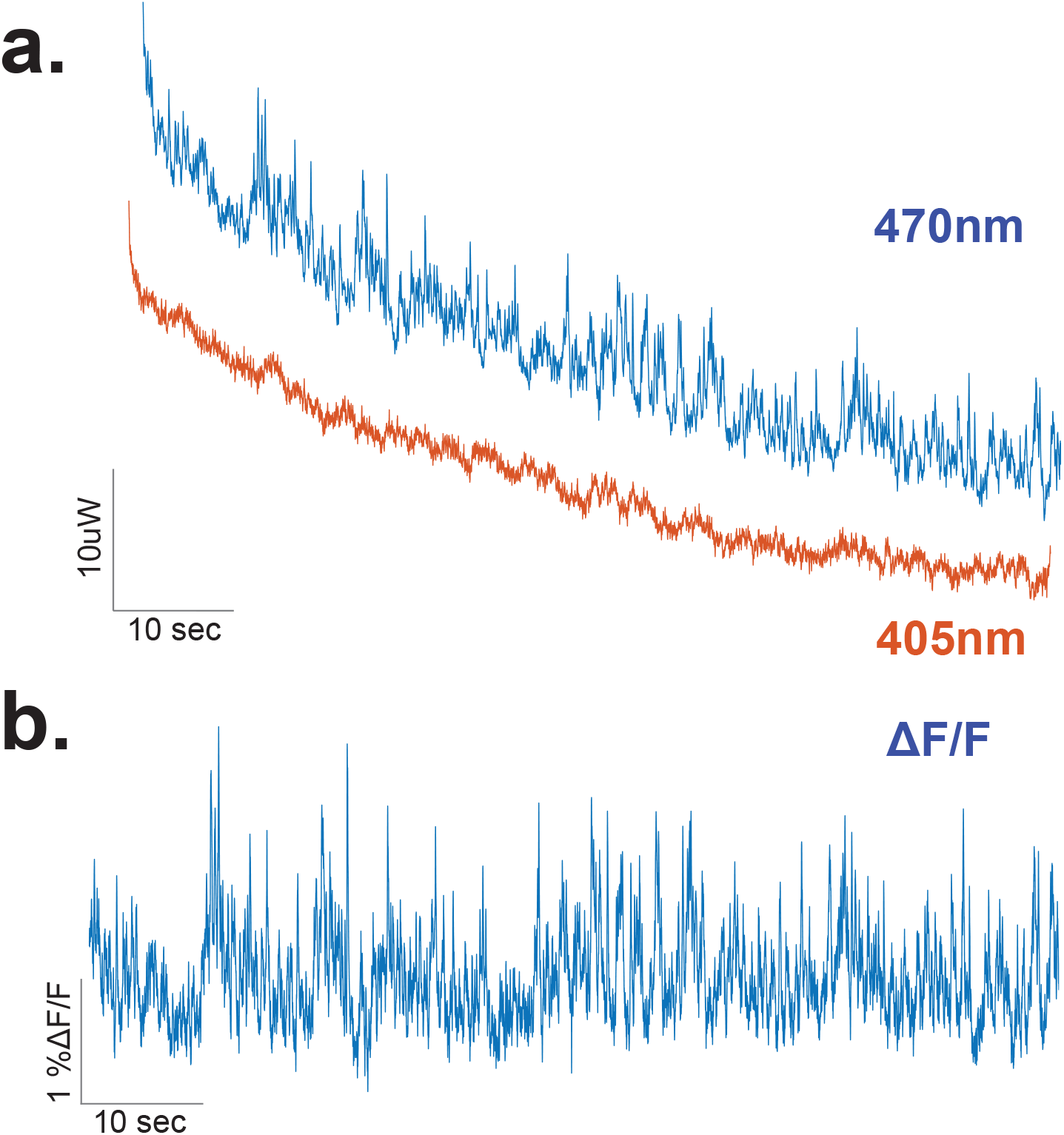
Representative dLight recording traces following optogenetic stimulation of NAc core terminals during training. **(a)** Representative traces for 470 nm excitation (dLight) and 405 nm excitation (isosbestic control) channels in an individual animal at baseline. **(b)** Representative ΔF/F trace showing dopamine transients in the nucleus accumbens core following optogenetic stimulation of the terminals via a red-shifted excitatory opsin (Chrimson).

**Figure S7.**
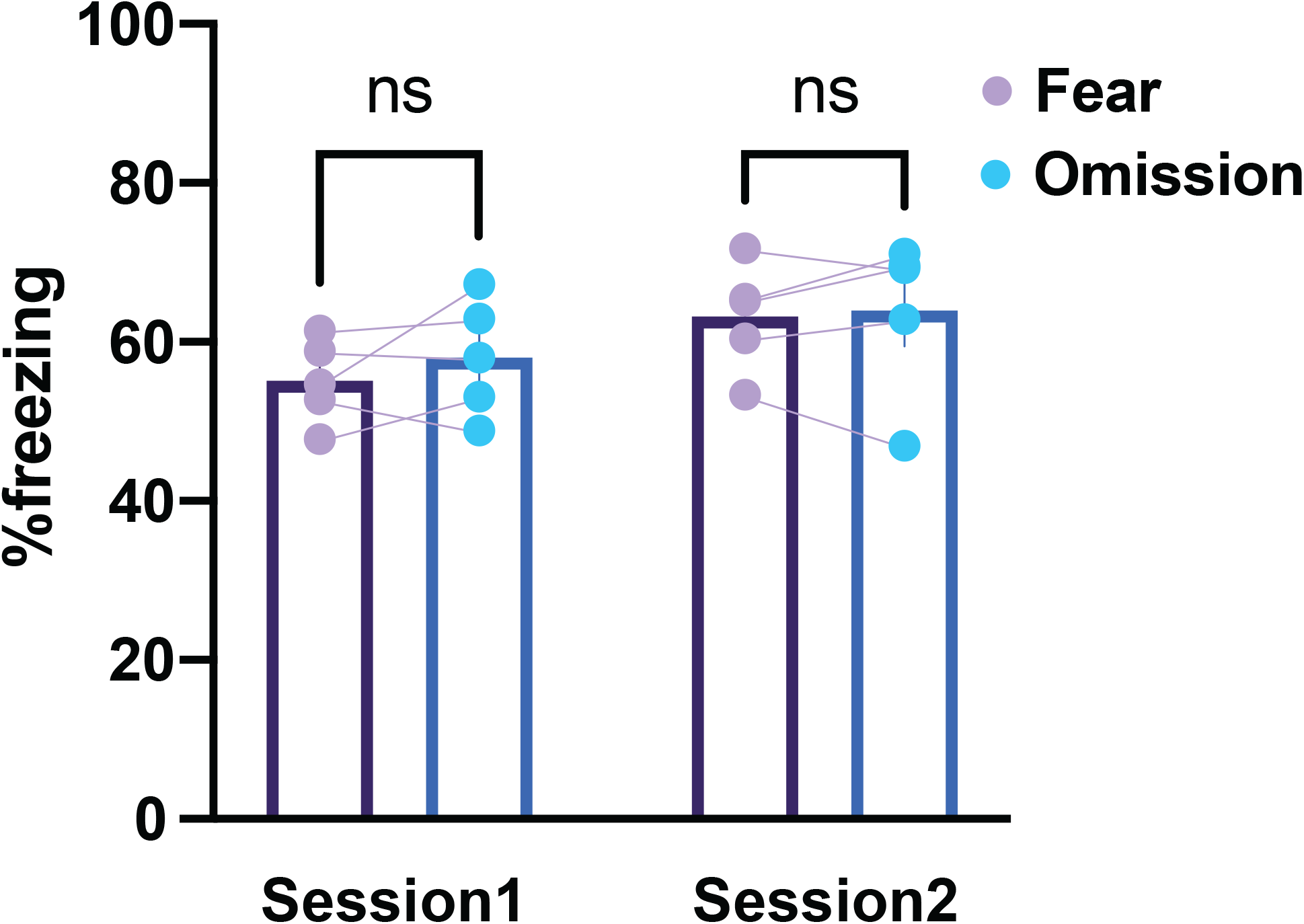
Mice failed to show robust inhibitory learning when the dopamine terminals are photostimulated during training. Averaged freezing response to the Fear+Omission cue did not differ from the freezing response to the Fear cue only trials during Session 1 (RM ANOVA Cue main effect *F*_(1,4)_= 0.67, *p*=0.4563; Sidak post-hoc p<0.01, n=5 mice) and Session 2 (Sidak post-hoc p<0.001, n=5 mice) of the aversive conditioned inhibition training. Data represented as mean ± S.E.M. ns = not significant.

**Figure S8.**
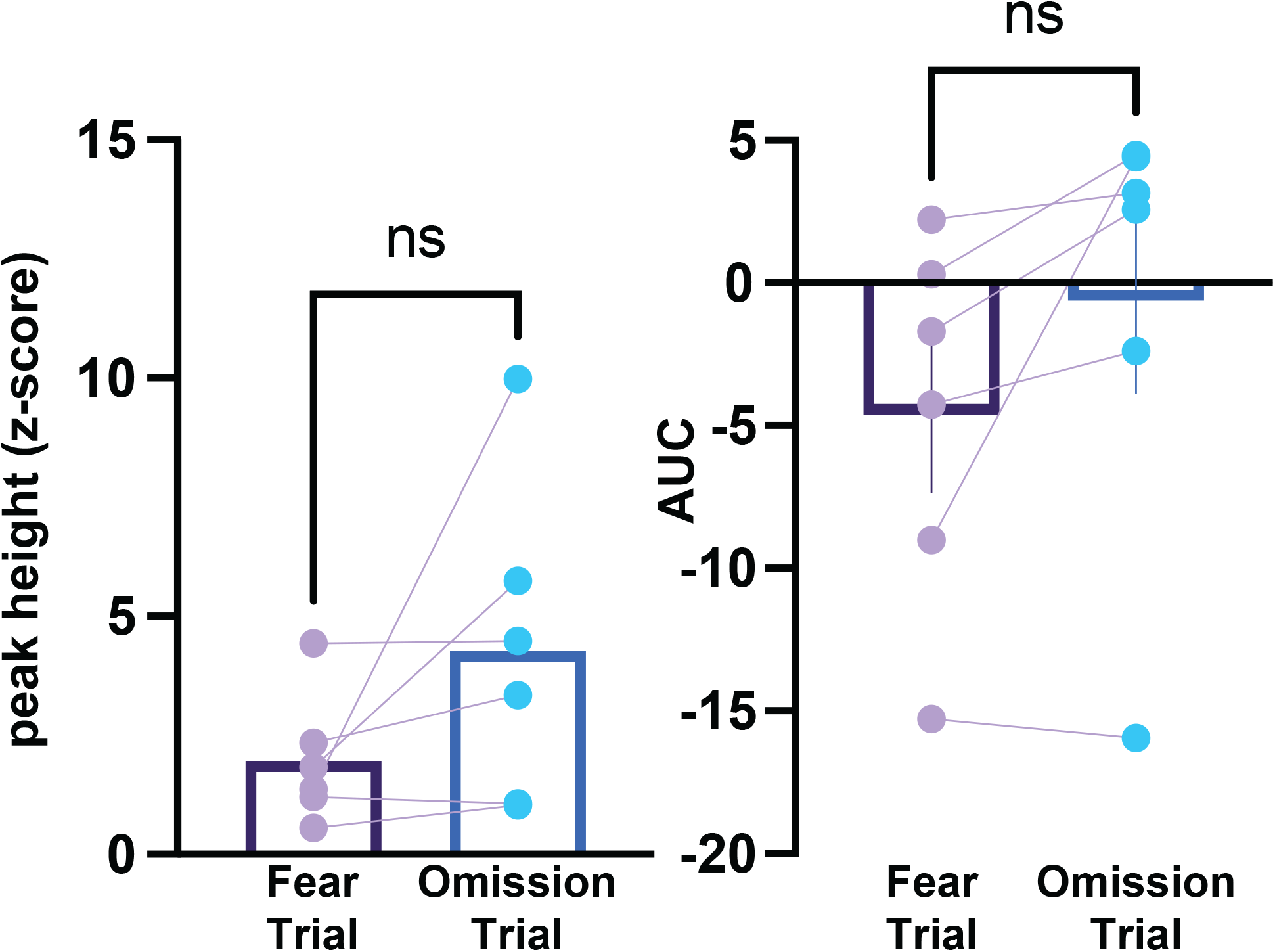
Averaged dopamine responses from individual animals did not differ at the time of the omitted shock following the Fear cue as compared to following the Fear+Omission cue. The peak-height (Paired t-test, *t*_4_=0.55, *p*=0.6114, n=5 mice) and the area under the curve (AUC; Paired t-test, *t*_4_=0.19, *p*=0.8514, n=5 mice) of the dopamine response to the expected but absent footshock at the offset of the Fear cue did not differ from the dopamine response at the offset of the Fear+Omission cue. Data represented as mean ± S.E.M. * *p* < 0.05.

**Figure S9.**
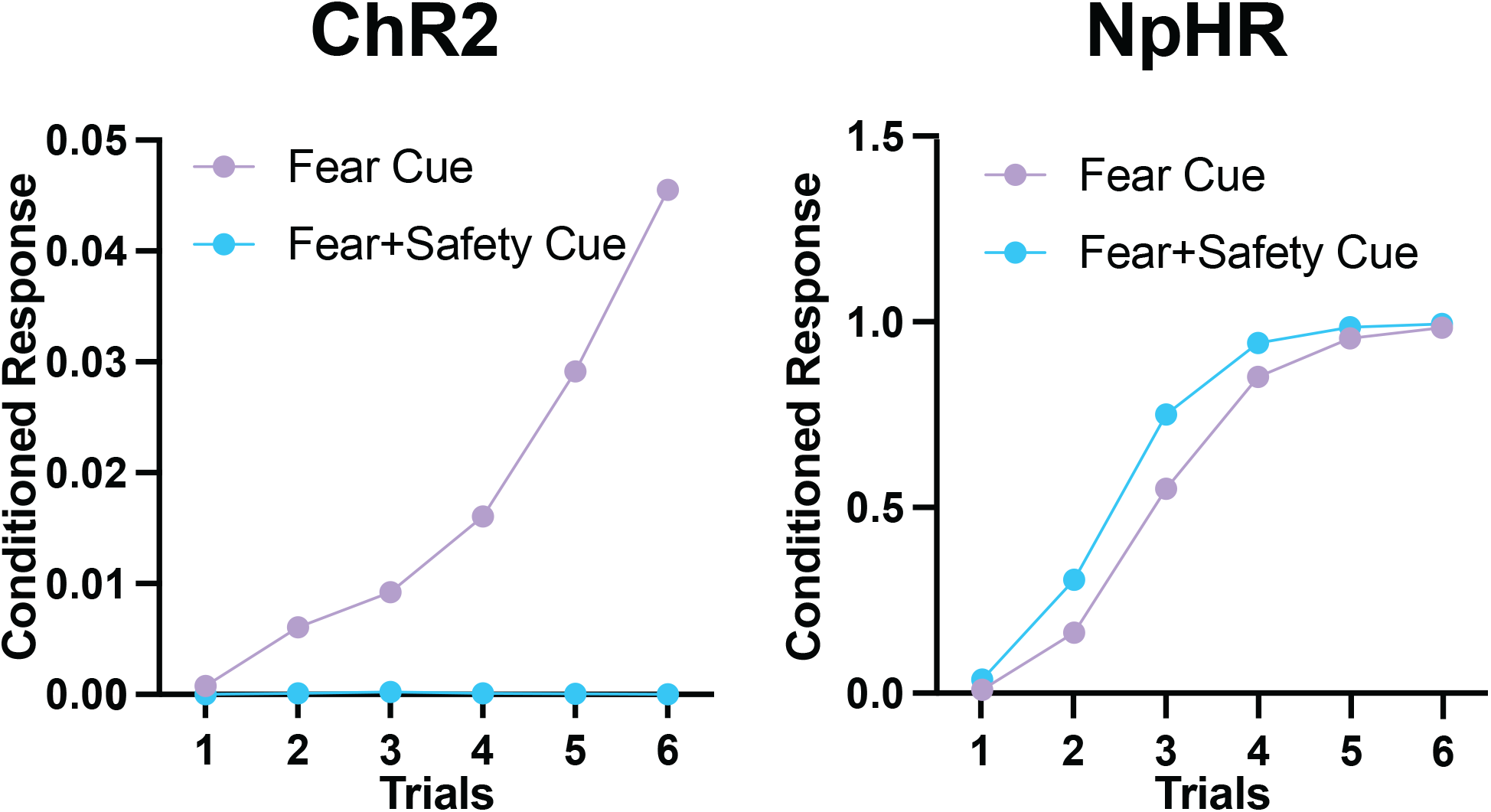
KCS model simulations: Artificially enhancing and suppressing the prediction error during conditioned inhibition. Our simulations of the KCS model showing that when the prediction error was set to a higher value (prediction error*2) to mimic the optogenetic stimulation of NAc dopamine (ChR2), opposite to our experimental data (see **Figure 2**), the KCS model learned conditioned inhibition faster. Conversely, when the prediction error was set to a negative value (prediction error*-1) to mimic optogenetic inhibition of dopamine terminals, the KCS model failed to learn conditioned inhibition rather than enhanced learning we observed in mice (see **Figure 2**).

**Figure S10.**
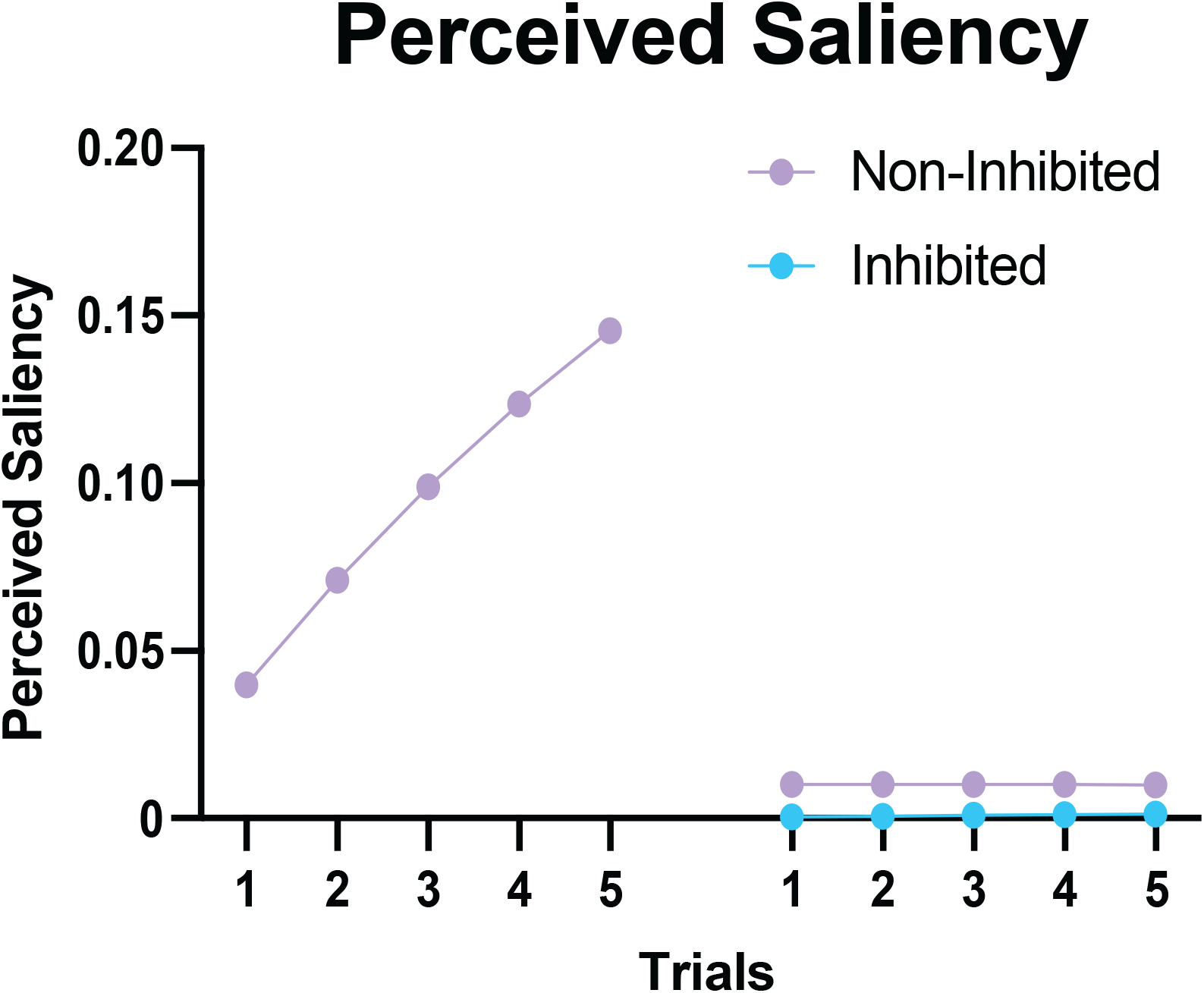
KCS model simulations: Perceived saliency following Inhibited vs. Non-Inhibited cue. KCS model simulations showing decreased perceived saliency during testing following the inhibition of the perceived saliency of the footshock outcome during training.

**Figure S11.**
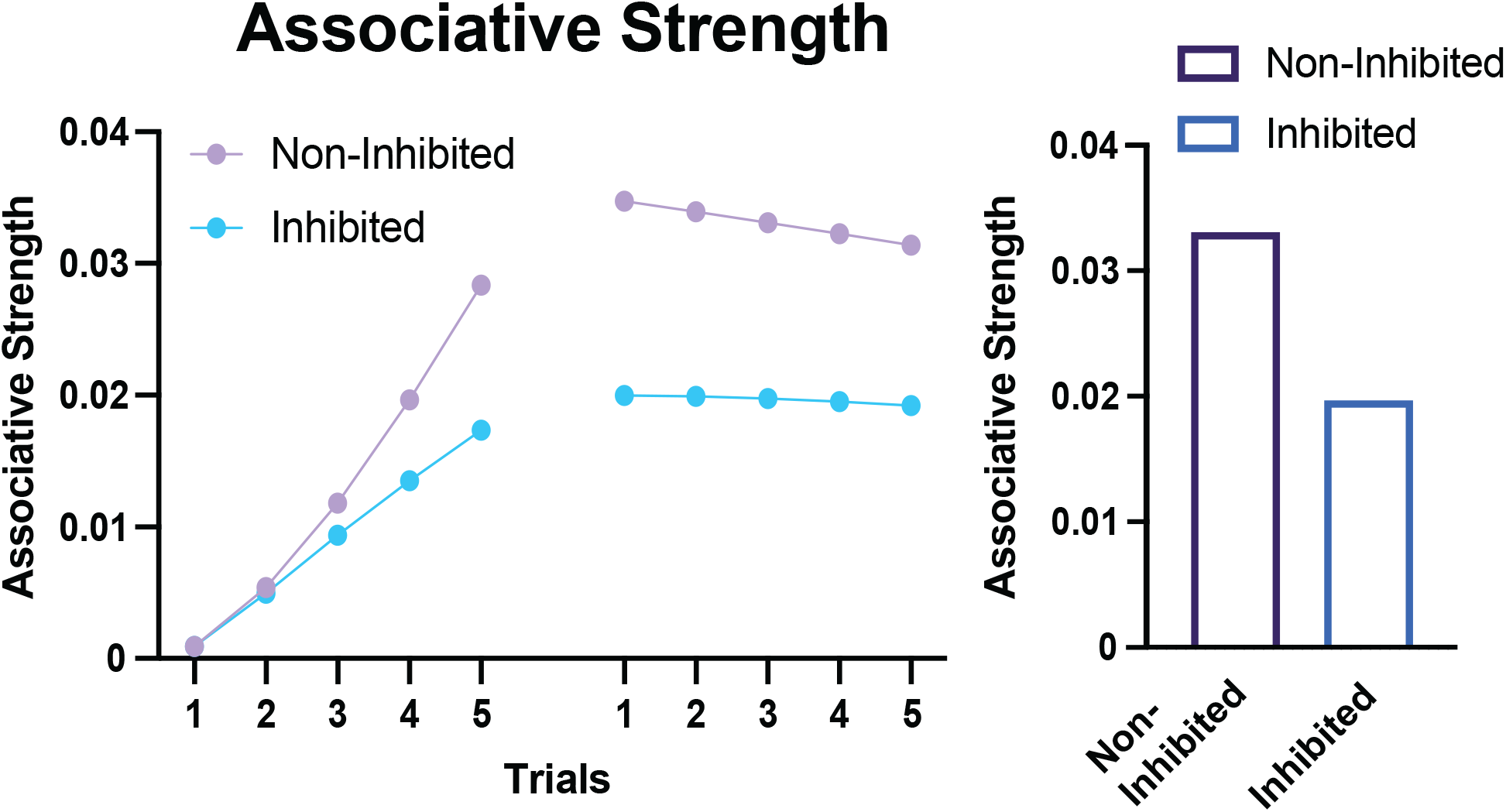
KCS model simulations: Associative strength following Inhibited vs. Non-Inhibited Cue. KCS model simulations showing weaker cue-outcome associations during testing following the inhibition of the perceived saliency of the footshock outcome during training.

**Figure S12.**
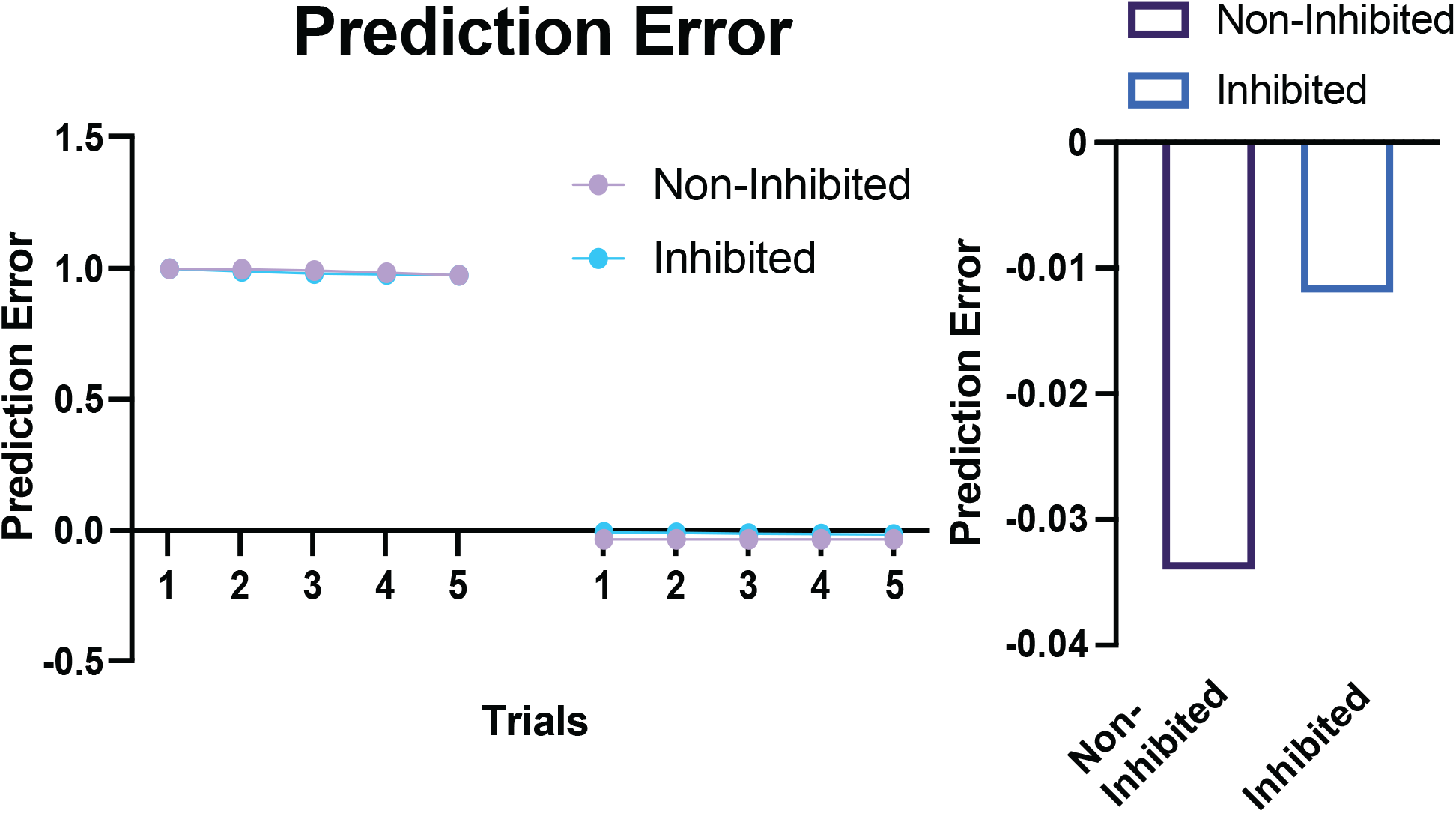
KCS model simulations: Prediction error following Inhibited vs. Non-Inhibited Cue. KCS model simulations showing reduced negative prediction error during testing following the inhibition of the perceived saliency of the footshock outcome during training.

**Figure S13.**
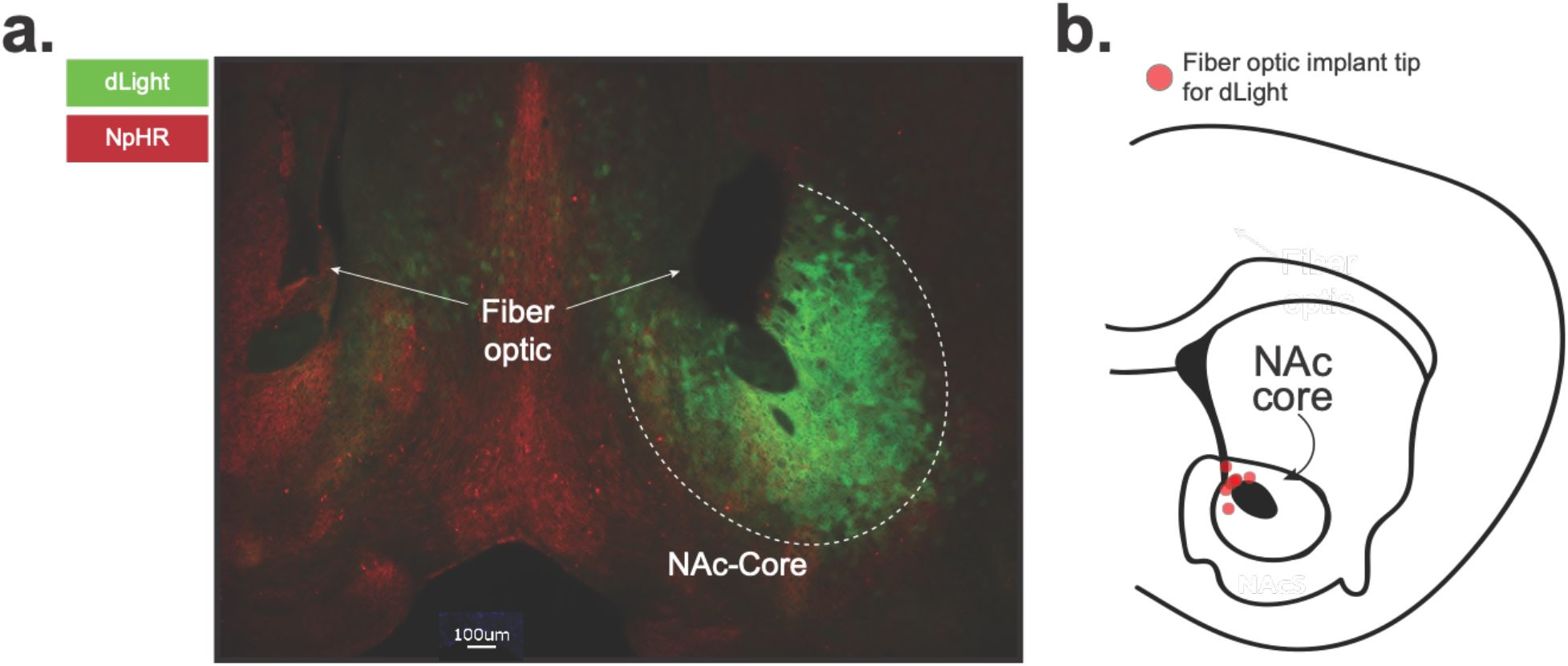
Expression of an Inhibitory opsin in dopamine terminals in conjunction with dLight in the NAc core. **(a)** A fiber optic cannula was placed directly above the NAc core in order to record dopamine transients via dLight with a blue LED. Concurrently terminals were inhibited via a yellow laser applied to dopamine terminals expressing halorhodopsin (AAV5.hSyn.eNpHR3.0.mCherry). Representative histology showing viral expression (green) of dLight and (red) halorhodopsin (NpHR) restricted to the NAc core and **(b)** schematic showing fiber optic placements (red) in experimental animals.

**Figure S14.**
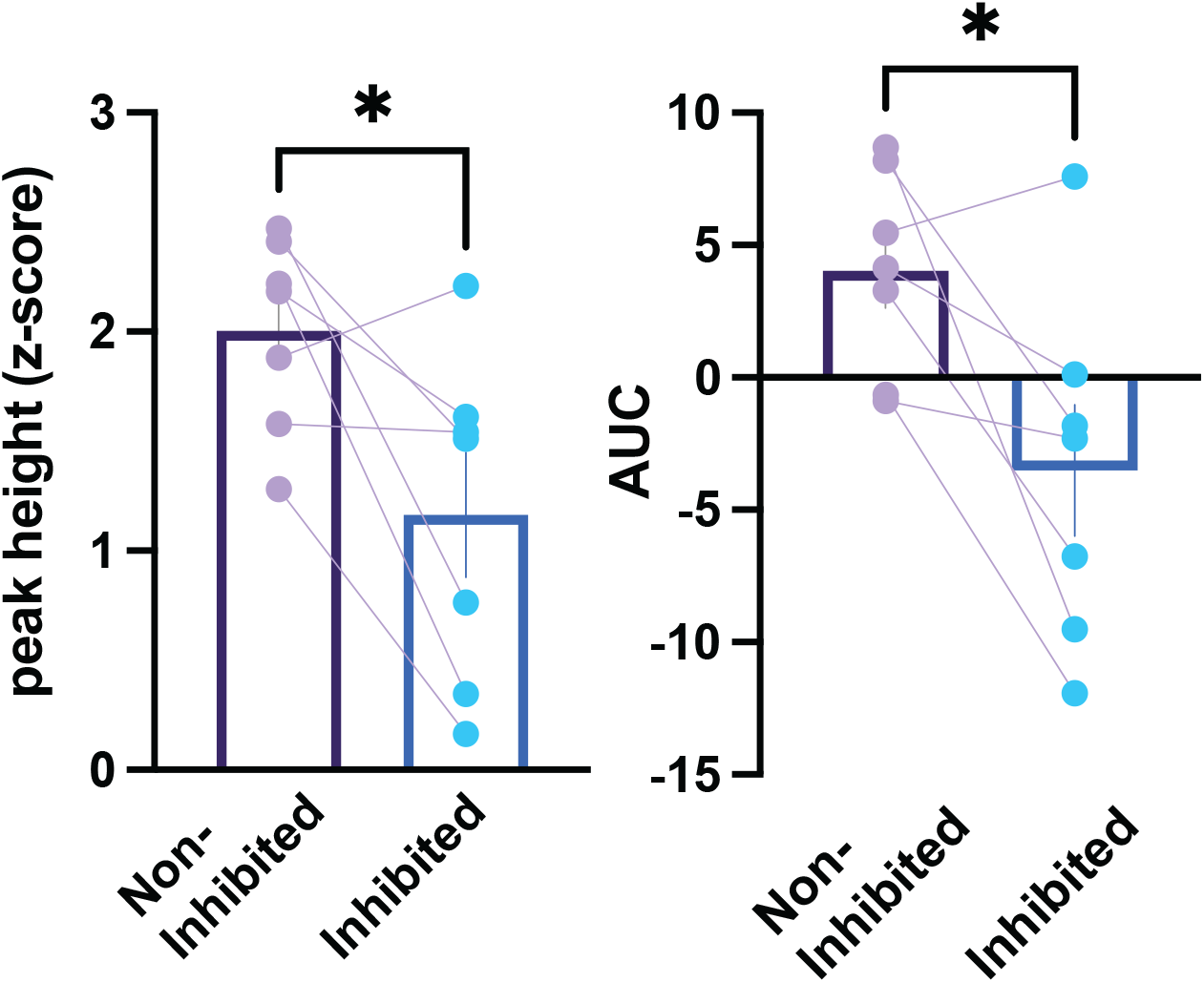
Averaged dopamine responses from individual animals show a stronger dopamine response during shock omission following the Non-inhibited cue. The peak-height (Paired t-test, *t*_6_=2.72, *p*=0.0342, n=7 mice) and the area under the curve (AUC; Paired t-test, *t*_6_=2.91, *p*=0.0270, n=7 mice) of the dopamine response to the expected but absent footshock at the offset of the Inhibited cue was smaller compared to the Non-inhibited cue offset. Data represented as mean ± S.E.M. * *p* < 0.05.

**Figure S15.**
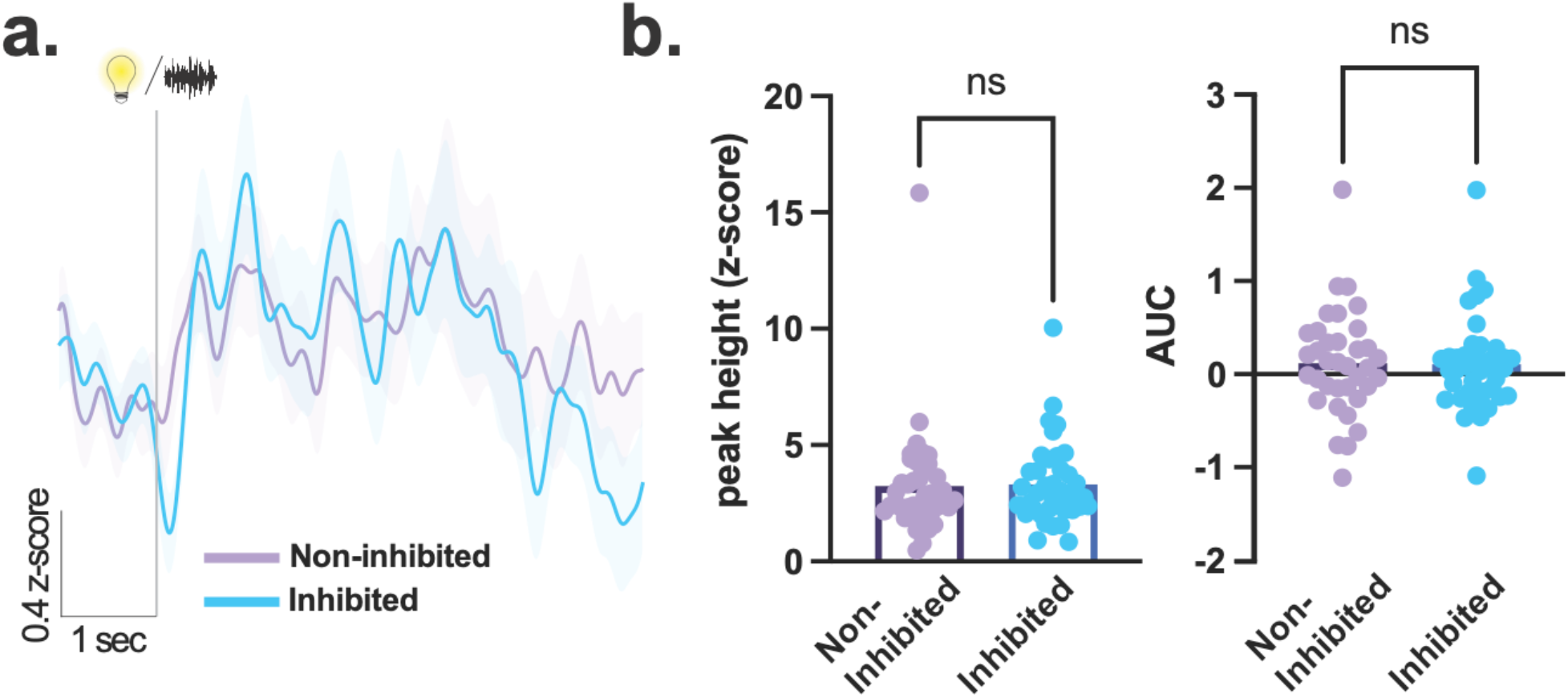
Inhibition of dopamine to the footshock during trainings did not affect dopamine responses to the predictive cues. **(a)** Dopamine response at the time of the Inhibited cue did not differ from the dopamine response to the Non-inhibited cue. **(b)** Peak height (Nested t-test, *t*_12_=0.22, *p*=0.8992, n=7 mice) and area under the curve (AUC; Nested t-test, *t*_12_=0.17, *p*=0.8644, n=7 mice) values of the dopamine response to the Inhibited and Non-inhibited cue did not differ. Data represented as mean ± S.E.M. ns = not significant.

## Notes

### Competing Interest Statement

The authors have declared no competing interest.

